# Nitric oxide modulates contrast suppression in a subset of mouse retinal ganglion cells

**DOI:** 10.1101/2023.12.23.572742

**Authors:** Dominic Gonschorek, Matías A. Goldin, Jonathan Oesterle, Tom Schwerd-Kleine, Ryan Arlinghaus, Zhijian Zhao, Timm Schubert, Olivier Marre, Thomas Euler

## Abstract

Neuromodulators have major influences on the regulation of neural circuit activity across the nervous system. Nitric oxide (NO) has been shown to be a prominent neuromodulator in many circuits and has been extensively studied in the retina. Here, it has been associated with the regulation of light adaptation, gain control, and gap junctional coupling, but its effect on the retinal output, specifically on the different types of retinal ganglion cells (RGCs), is still poorly understood. In this study, we used two-photon Ca^2+^imaging and multi-electrode array (MEA) recordings to measure light-evoked activity of RGCs in the ganglion cell layer in the *ex vivo* mouse retina. This approach allowed us to investigate the neuromodulatory effects of NO on a cell type-level. Our findings reveal that NO selectively modulates the suppression of temporal responses in a distinct subset of contrast-suppressed RGC types, increasing their activity without altering the spatial properties of their receptive fields. Given that under photopic conditions, NO release is triggered by quick changes in light levels, we propose that these RGC types signal fast contrast changes to higher visual regions. Remarkably, we found that about one-third of the RGC types, recorded using two-photon Ca^2+^imaging, exhibited consistent, cell type-specific adaptational response changes throughout an experiment, independent of NO. By employing a sequential-recording paradigm, we could disentangle those additional adaptational response changes from drug-induced modulations. Taken together, our research highlights the selective neuromodulatory effects of NO on RGCs and emphasizes the need of considering non-pharmacological activity changes, like adaptation, in such study designs.

## Introduction

The retina can be considered as a visual signal processor with a straightforward functional architecture (1–3). The photoreceptors convert the stream of photons from the environment into an electrical signal, which is relayed downstream by the bipolar cells (BCs) to the retinal ganglion cells (RGCs), the tissue’s output neurons. Along this vertical pathway, the visual signal is shaped by two lateral, inhibitory cell classes: horizontal cells in the outer and amacrine cells (ACs) in the inner retina. The resulting intricate networks allow sophisticated computations, reflected in the >40 output channels that relay diverse visual feature representations as spike trains to higher brain regions (4–6).

Over the past decades, the retinal synaptic networks have been intensively studied, which greatly broadened our understanding of early visual signal processing (7–12). Specifically, the availability of connectome data for large parts of the retina helped in unprecedented ways, as can be seen from studies, for example, investigating direction-selective circuits and their precise wiring regarding starburst ACs and direction-selective RGCs (e.g., (13, 14)).

What is not well-captured by connectomic approaches are ‘wireless’ interactions mediated by neuromodulators, which comprise a broad variety of very different small molecules, including monoamines (e.g., dopamine (15, 16), histamine (17, 18), serotonin (19)), endocannabinoids (20– 22), gasotransmitters such as nitric oxide (NO) (23, 24)), as well as a variety of neuropeptides (e.g., neuropeptide Y (25, 26)). Only few of the >20 neuromodulators (27) found in the retina are released from centrifugal fibers (e.g., histamine and serotonin (17–19, 28)), whereas most of them are released in addition to GABA or glycine by ACs (27, 29– 32). Neuromodulators have long been implicated in adapting the retina to different contextual states necessary to robustly perform in a highly dynamic visual environment (e.g., (33–36)). For instance, dopamine (DA) is released by a distinct type of AC (37–40), and has been shown to regulate light adaptation (33, 41, 42) via several cellular mechanisms, such as modulation of gap junctional coupling (34, 43–45) and intrinsic conductances (46, 47). More recently, histamine was proposed to shape the retinal code in a top-down modulatory manner related to the animal’s arousal state (17, 28). For several neuromodulators, the retinal release sites and receptors are known and their effects on cellular properties have been described, however, a comprehensive, function-oriented, and cell-type view of these neuromodulators’ functional implications and contributions to visual signal processing is only slowly emerging (e.g., (17, 48)).

One of the better-studied neuromodulators in the retina is NO. Neuronal nitric oxide synthase (nNOS) is considered the dominant enzyme producing NO relevant for retinal signal processing, and — depending on species — was shown to be present in different retinal cell classes (49–57). In the mouse retina, nNOS was mainly found in specific AC types (36, 58–60). A few years ago, Jacoby et al. (36) demonstrated that one of those types (nNOS-2 AC) controls the light-dependent release of NO in the inner retina. NO can function via two main pathways (61): (*i*) it can bind to the NO guanylate cyclase (NO-GC; also referred to as soluble guanylate cyclase (sGC)) receptor, triggering the production of the second messenger cyclic guanosine monophosphate (cGMP), which binds to downstream targets (55, 62, 63), and (*ii*) via S-nitrosylation (cGMP-independent) by directly modifying certain receptor and channel proteins (64, 65).

The effects of NO have been primarily linked with light adaptation and the transition between scotopic and photopic signaling pathways via several mechanisms, including uncoupling the gap junctions between AII ACs and On-cone BCs (66), increasing the gain of On-cone BCs in response to dim light (67, 68), and modulating the gain of Off-cone BCs (69, 70). At the retina’s output level, increasing NO was found to decrease On- and Off-responses in RGCs (71). Notably, genetically knocking out nNOS also led to a reduced light sensitivity in RGCs (72), which was also shown by inhibiting nNOS under light-adapted conditions (73). Additionally, NO has been shown to modulate RGC responses via cGMP by altering their cGMP-gated conductances (74, 75). Taken together, NO can act on different levels and via different pathways in the retina. However, the function of NO at the cell type-level and its role in early visual processing is far from understood.

Here, we systematically studied the functional role of NO and its effects on all retinal output channels in the *ex vivo* mouse retina. Surprisingly, we observed highly reproducible, cell type-specific changes in the light responses of some RGCs already in the absence of pharmacological manipulation. To account for these adaptational response changes, we developed a recording paradigm to sequentially measure RGC responses under control and drug conditions. We found that NO had a highly selective effect on a distinct set of three RGC types characterized by their suppressed response to temporal contrast, where NO strongly and differentially reduced this suppression, as well as caused them to respond faster. Yet, NO had no discernible effects on their spatial receptive field properties. Together, our data suggest that NO modulates the visual feature representation of the retinal output in a highly selective and type-specific manner.

## Results

To investigate NO effects systematically on the retinal output, we performed population imaging from somata in the ganglion cell layer (GCL) of *ex vivo* mouse retina electroporated with the synthetic Ca^2+^ indicator Oregon Green BAPTA-1 (Fig. 1a-c; Methods). In addition, we recorded a complementary dataset using multi-electrode array (MEA) recordings. To identify the different types of RGCs and detect potential NO-induced response changes, we presented a set of visual stimuli, including full-field chirps, moving bars (Fig. 1d), and binary dense noise (5) (see Methods).

**Fig. 1.**
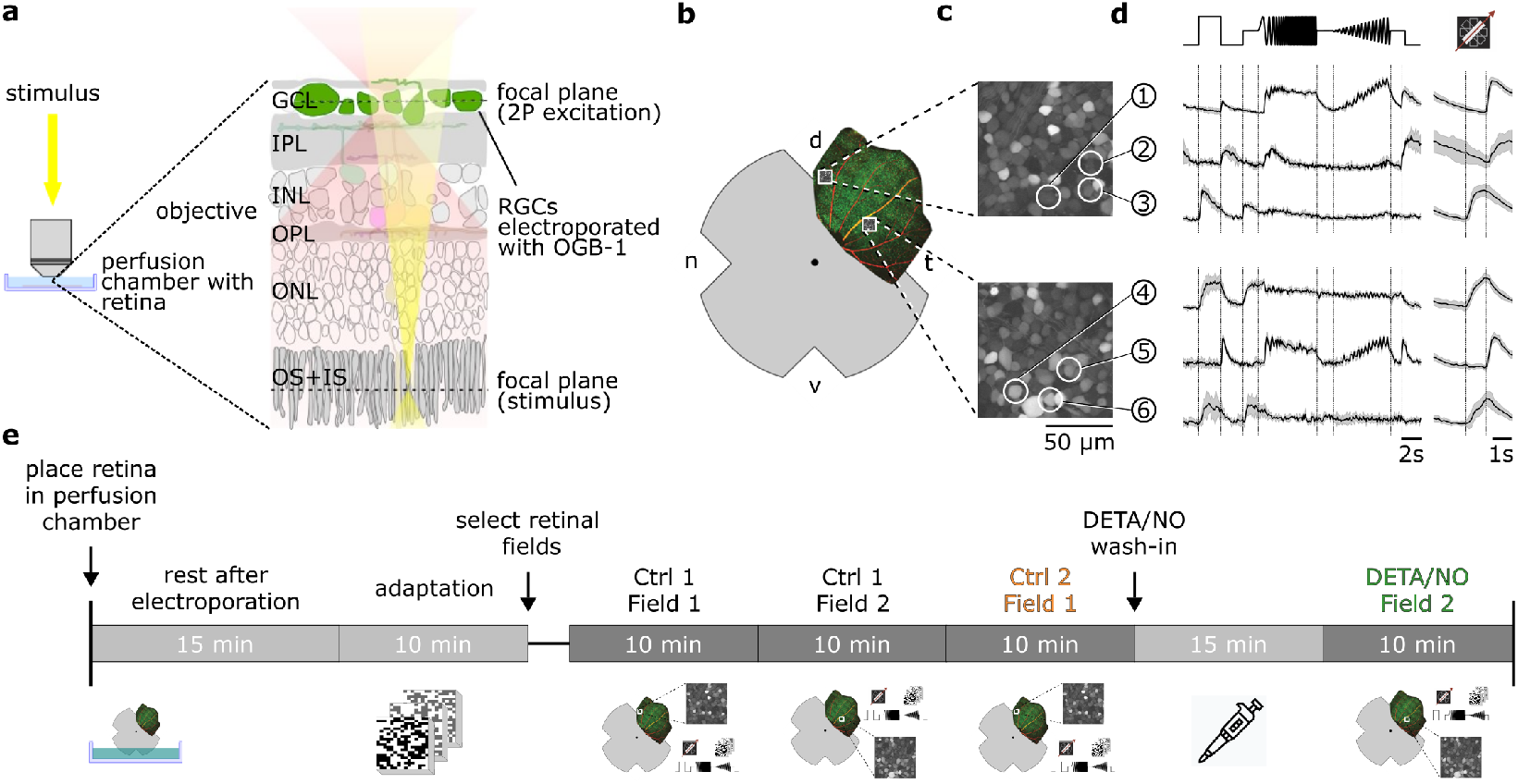
Overview of the experimental setup and recording procedure. (**a**) Two-photon imaging of ganglion cell layer (GCL) somata in the whole-mounted *ex vivo* mouse retina. (**b**) Schematic *ex vivo* whole-mounted retina (dot marks optic disc; d, dorsal; t, temporal; v, ventral; n, nasal). (**c**) Two representative recording fields from (b) showing GCL somata loaded with Ca^2+^ indicator OGB-1 (Methods). (**d**) Representative Ca^2+^ activity from cells in the GCL (white circles in (c)) in response to chirp (left) and moving bar stimulus (right) (black, mean; gray, s.d.). (**e**) Timeline of experimental procedure; for details, see text. IPL, inner plexiform layer; INL, inner nuclear layer; OPL, outer plexiform layer; ONL, outer nuclear layer; OS+IS, outer and inner segments; DETA/NO, nitric oxide (NO) donor.

### A protocol for sequential control/drug recordings

Previous studies have shown that retinal responses recorded with two-photon Ca^2+^ imaging can be systematically affected by experimental factors, such as excitation laser-induced activity, photoreceptor bleaching, and temporal filtering due to Ca^2+^ buffering by the fluorescent indicator (76–78). These changes can be summarized by the umbrella term ‘batch effects’ (a term coined in the molecular genetics field), which can confound the biological signal and potentially cause erroneous interpretations of the data (78, 79). Such batch effects may play a role when, as in our study, data are recorded in a sequential manner to infer possible drug effects.

Because we wanted to detect potentially subtle NO effects, we devised a protocol to make experiments as comparable as possible (Fig. 1e). After placing the tissue into the recording chamber, it was allowed to recover from the electroporation for 15 min, before we light-adapted the retina for 10 min by presenting the dense noise stimulus. We then selected retinal recording fields, each of which was recorded twice for the complete stimulus set in an interlaced manner (Fig. 1e). Here, we made sure that the fields were uniformly labeled with the Ca^2+^ indicator and responsive to light stimuli. The first field was recorded twice without perturbation (*control-control*). For the next field, we added the drug to the perfusion medium and incubated the tissue for ∼15 min, before recording the field for the second time (*control-drug*). For the last 5-10 min of the wash-in time, we presented the noise stimulus to preserve the light adaptation level. Note that the time between the 1^st^ and 2^nd^ recording of field 2 was ∼15 min longer (the wash-in time) than that of field 1 (see Discussion). For the Ca^2+^ data, we decided against recording also after wash-out, because response quality decreased for the 3^rd^ scan of the same field, likely due to bleaching of fluorescent indicator and photopigment. However, we did include wash-out in the MEA-dataset (see below).

Our sequential-recording protocol yielded paired-data at the cell-level, allowing us to track if and how each cell’s responses changed under the drug condition. Using this protocol, we recorded the following datasets: (*i*) a control-dataset to test response stability, i.e., Ctrl 1 and Ctrl 2, (*ii*) a strychnine-dataset to test the protocol for a drug with well-described effects, i.e., Ctrl 1 and Strychnine (1 μM), and *(iii*) a NO-dataset to infer NO-induced response changes, i.e., Ctrl 1 and NO (DETA/NO; 100 μM). The control-dataset was leveraged to reveal NO-induced effects on the background of potential non-specific response changes throughout the experiment. For the following analyses, we used 3,975 RGCs (n_Ctrl_=1,701; n_NO_=1,838, n_Strychnine_=436) that fulfilled our response quality filtering (see Methods).

### Identifying functional RGC types using a classifier

The mouse retina contains more than 40 RGC types (5, 6). As we wanted to investigate if the tested drugs differentially affect the different retinal output channels, we applied an RGC type classifier (Fig. 2a) (80), which had been trained and validated on a previously published RGC Ca^2+^ imaging dataset (5). The classifier predicts a GCL cell’s functional type based on soma size and the responses to chirp and moving bar stimuli (see Methods). The classifier also distinguishes between RGCs and displaced ACs. Here, we focused our analysis on the RGCs. To match the conditions under which the classifier’s training data was acquired as closely as possible, we predicted types based on the responses from the first control recording (Ctrl 1) and the cells retained their type over the course of their experiment (no re-typing). To minimize classification uncertainty, we additionally discarded cells with low confidence scores (*<* 0.25, see Methods for details).

**Fig. 2.**
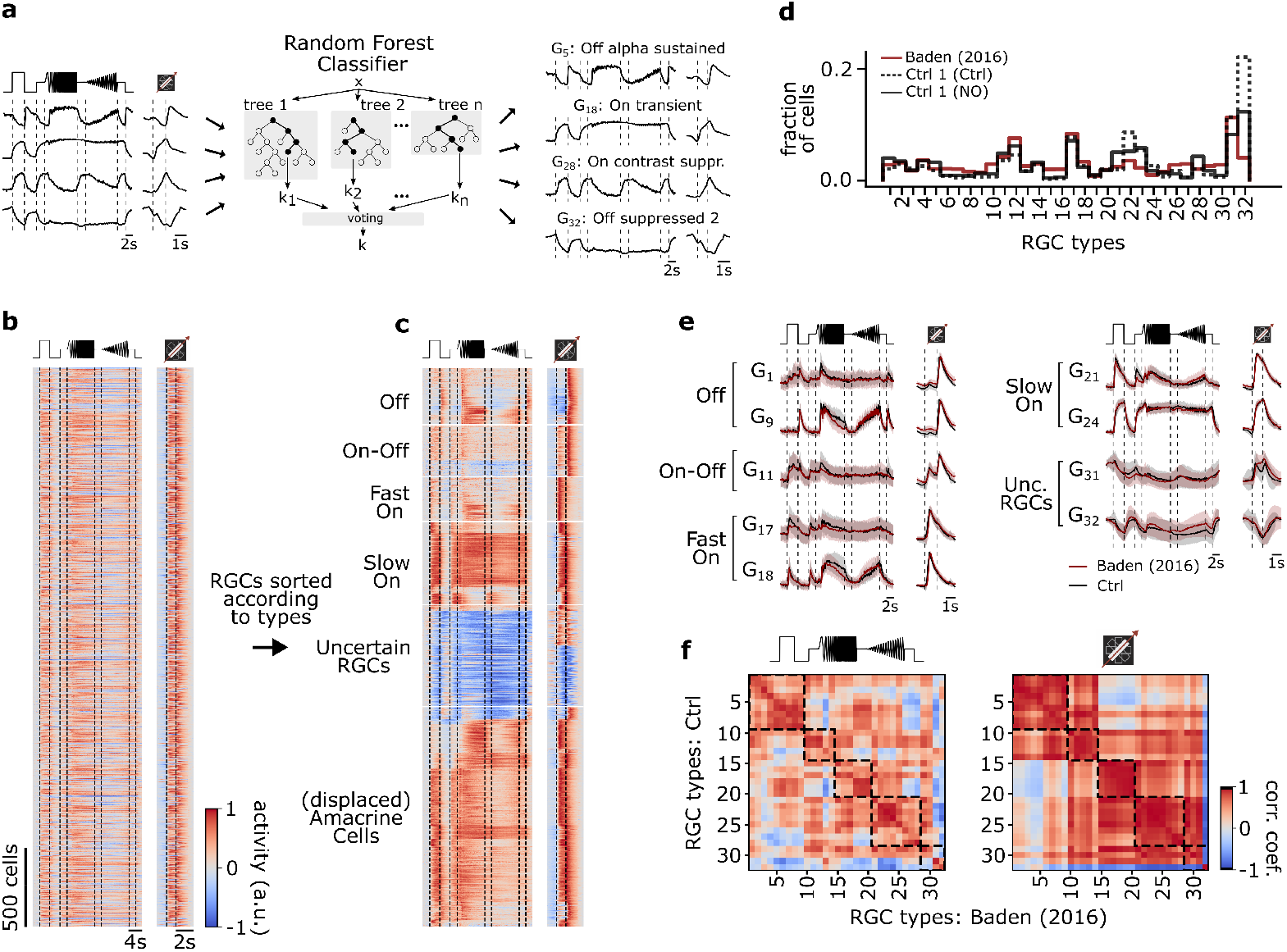
Functional classification of mouse RGC types. (**a**) Illustration of the random forest classifier (RFC) to predict cell type labels for Ctrl 1 of both datasets. For each cell, Ca^2+^ responses to chirp and moving bar, soma size, and p-value of permutation test for direction selectivity (left) constitute the input to the RFC (center) to predict a cell type label, i.e., a type GX (right). For details, see Methods and (80). (**b**) Pooled heat map of unsorted cell responses to chirp and moving bar stimulus from both Ctrl 1 datasets after quality filtering (QIMB>0.6 or QIchirp>0.45, and classifier confidence score *≥* 0.25). The color bar indicates normalized response activity. (**c**) Heat map from (b), but sorted according to their assigned type. (**d**) Distribution of RGC types predicted by the RFC classifier for both Ctrl 1 of the control(Ctrl 1 (Ctrl); solid black), of DETA/NO-dataset (Ctrl 1 (NO); dotted black), and for the dataset from Baden et al. (5) (red). (**e**) Representative RGC type response averages to chirp and moving bar (Ctrl, black; training dataset, red). (**f**) Correlation matrix of type mean responses per RGC type between Ctrl and Baden et al. (5) dataset for chirp (left) and moving bar (right). Dashed boxes indicate functional groups (Off, On-Off, Fast On, Slow On, and Uncertain RGCs; see (5)). The color bar indicates the Pearson correlation coefficient.

We found that the distributions of the predicted RGC types in our datasets matched that of the earlier dataset quite well (Fig. 2b-d). Also, the predicted mean traces for the individual RGC types were very similar to those in Baden et al. (5) (Fig. 2e), as indicated by the high correlations of their chirp and moving bar responses (Fig. 2f left and right, respectively). That the moving bar responses were more strongly correlated than the chirp responses is likely due to the lower complexity and shorter duration of the former stimulus. Nonetheless, we found the RGC classification overall to be robust and comparable to the original dataset (Fig. S1a,b).

### Testing the recording protocol

In the mouse retina, glycine is released by small-field ACs, which relay inhibition vertically across the inner plexiform layer (IPL) and thus, are involved in cross-over inhibition (32, 81). Blocking cross-over inhibition between the On- and Off-pathways is therefore expected to have effects on many RGC circuits, which is why we chose the glycine receptor antagonist strychnine to test our recording protocol. Specifically, we focused on strychnine unmasking responses to the other stimulus polarity (82). Indeed, we found that strychnine revealed additional On-response components in Off (e.g., G_1_, G_2_, G_4_, G_6_) and On-Off RGCs (e.g., G_11_, G_12_), as can be seen, for instance, in their leading-edge response to the moving bar (Fig. S2). In On RGCs, we did not detect additional (Off) response components. Instead, some On RGCs exhibited slightly more sustained responses to light increments (e.g., G_18_, G_20_, G_22_). Together, the strychnine-dataset demonstrates that we can resolve drug-related effects on light responses at the RGC type-level.

### Certain RGC types display adaptational response changes

To test if our recording conditions were stable and to exclude major batch effects, we first compared the responses of the control-datasets (Ctrl 1 vs. Ctrl 2). To this end, we computed the difference between the Ctrl 1 and Ctrl 2 mean responses (ΔCtrl: ΔR_(Ctrl2-Ctrl1)_) to chirp and moving bar stimuli for each cell of every RGC type. This allowed us to quantify if and how the responses changed over the time course of an experiment (cf. protocol in Fig. 1e). Here, we only considered RGC types with >10 sequentially recorded cells (21/32).

Surprisingly, while the majority of RGC types featured stable responses (e.g., G_1_, G_21_; Fig. 3a), a substantial number of RGC types (9/21) changed their responses to chirp and/or moving bar stimuli in the absence of any pharmacological perturbation in a highly reproducible manner (Fig. S3a,b). For instance, for Ctrl 2, G_23_ showed reduced responses, whereas G_31_ showed an increased response activity (Fig. 3b). Interestingly, cells assigned to the functional groups of ‘Off’ RGC types displayed stable responses, whereas ‘On-Off’, ‘Fast On’, ‘Slow On’, and ‘Uncertain RGCs’ included types with changing responses (50% (2/4), 34% (1/3), 67% (4/6), and 67% (2/3), respectively). This diversity argues against a systematic effect (such as, e.g., general run-down) and for a cell type-specific phenomenon, which in the following we refer to as ‘adaptational response changes’.

**Fig. 3.**
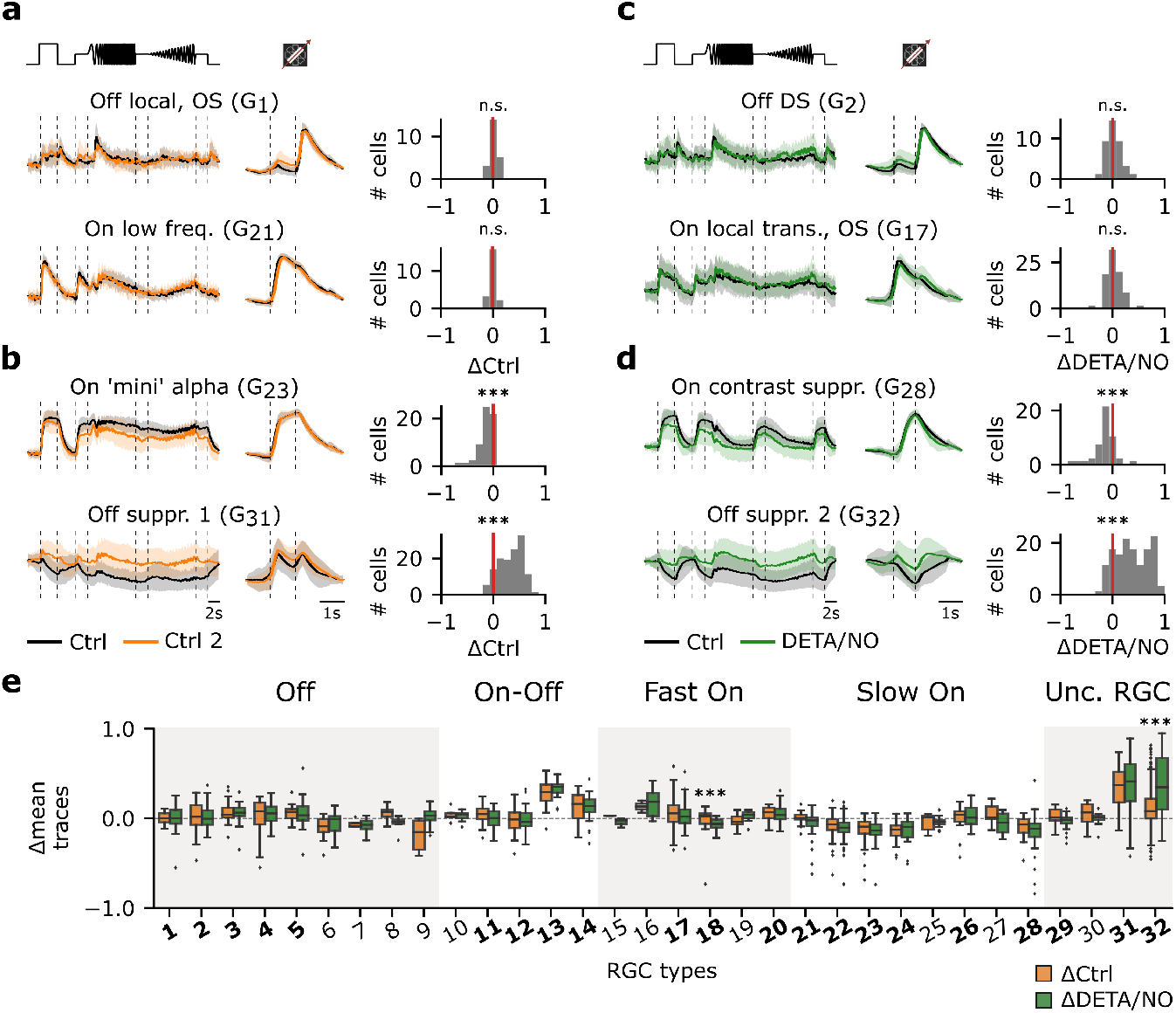
Certain RGC types are affected by adaptational and/or NO-induced effects, while others are unaffected. (**a**) Left: Two representative mean Ca^2+^ responses of sequentially recorded RGC types showing no differences between Ctrl 1 (black) and Ctrl 2 (orange) (top: G_1_; bottom: G_21_). Right: Corresponding histograms displaying the differences between the average traces of the sequentially recorded cell of the respective cell types. Zero indicates no difference between the response of the same cell across both recordings, whereas negative values indicate a decreased and positive values an increased activity. n.s.: not significant; ***: p<0.001; One-sample T-test. (**b**) Two representative RGC types that show decreased (top: G_23_) and increased (bottom: G_31_) response activity during Ctrl 2. n.s.: not significant; ***: p<0.001; One-sample T-test. (**c**) As in (a), but between sequentially recorded Ctrl 1 (black) and DETA/NO (green) (top: G_2_; bottom: G_17_). n.s.: not significant; ***: p<0.001; One-sample T-test. (**d**) As (c), but showing two cell types that display a decreased (top: G_28_) and increased (bottom: G_32_) activity when perfused with DETA/NO. n.s.: not significant; ***: p<0.001; One-sample T-test. (**e**) Box plots of trace differences of all sequentially recorded cells of all RGC types from control(ΔCtrl: ΔR_Ctrl2-Ctrl1_; orange) and NO-dataset (ΔDETA/NO: ΔR_NO-Ctrl1_; green). Bold numbers indicate RGC types with >10 sequentially recorded cells per dataset and condition. Dashed line shows zero baselines, i.e., no difference between traces. Diamond symbols represent outliers. Gray and white background blocks summarize the larger functional groups for better visualization (Off, On-Off, Fast On, Slow On, Uncertain RGCs). ***: p<0.001; Mann-Whitney U-Test.

### NO affects retinal output in a highly type-specific manner

Next, we investigated the effects of NO on the RGC responses. As with the control-dataset, we computed the cell-wise response differences between Ctrl 1 and NO responses (ΔDETA/NO: ΔR_NO-Ctrl1_). Similar to the control-dataset, the majority of RGC types displayed stable responses (e.g., G_2_, G_17_; Fig. 3c), while ∼43% changed their responses significantly (e.g., G_28_, G_32_; Fig. 3d) following the NO perfusion. We found that the percentage of changing types per functional group was similar to that in the control-dataset: ‘Off’ (0% (0/5)), ‘On-Off’ (50% (2/4)), ‘Fast On’ (34% (1/3)), ‘Slow On’ (66% (4/6)), and ‘Uncertain RGCs’ (66% (2/3)). This raised the question if the observed changes in the NO-dataset indeed reflected NO-induced modulations or mostly adaptational response changes (as observed in the control-dataset). We, therefore, tested for each RGC type if the response changes observed for control (ΔCtrl: ΔR_Ctrl2-Ctrl1_) and NO (ΔDETA/NO: ΔR_NO-Ctrl1_) were significantly different (Fig. 3e). To our surprise, this was only the case for two types: (1) G_32_ (‘Off suppressed 2’) RGC, which is characterized by a high baseline activity that is strongly suppressed below baseline during light increments and displays increased activity during light decrements, and (2) G_18_, which is considered as being a ‘Fast On’ type having a response to light increments with a fast response kinetic. This suggests highly type-selective NO effects — at least for temporal responses to chirp and moving bar stimuli.

Next, we leveraged the control-dataset to disentangle NO-induced from adaptational effects at the level of the response features. To this end, we subdivided the chirp stimulus into six feature segments ((1) On, (2) Off, (3) low frequency, (4) high frequency, (5) low contrast, (6) high contrast), and the moving bar into two ((7) On, (8) Off) (Fig. 4c). Then, for every cell type and every response feature, we computed the difference of the mean responses between the first and second recording separately for the control (Fig. 4a; left panel) and NO-dataset (Fig. 4a; middle panel). To isolate NO-induced effects, we computed the differences (ΔR_NO-Ctrl1_ ΔR_Ctrl2-Ctrl1_; Fig. 4a; right panel), based on the assumption that the adaptational component of the changes would be similar for both datasets. Through this analysis, we found three response behaviors across the RGC types: (1) not NO-affected/stable, (2) showing adaptational cell type-specific changes, and (3) NO-affected (Fig. 4b). As before, we found that G_32_ is strongly affected by NO; it showed barely any adaptational response changes, yet its activity increased during NO application (Fig. 4b; (3)). This increase in activity was statistically significant for the chirp’s On (1) and Off steps (2) and during both the frequency and contrast modulations ((3)-(6)), as well as for the moving bars trailing edge (8) (for tests, see Fig. 4 legend), suggesting that NO reduces the cell’s inhibition by temporal contrast.

**Fig. 4.**
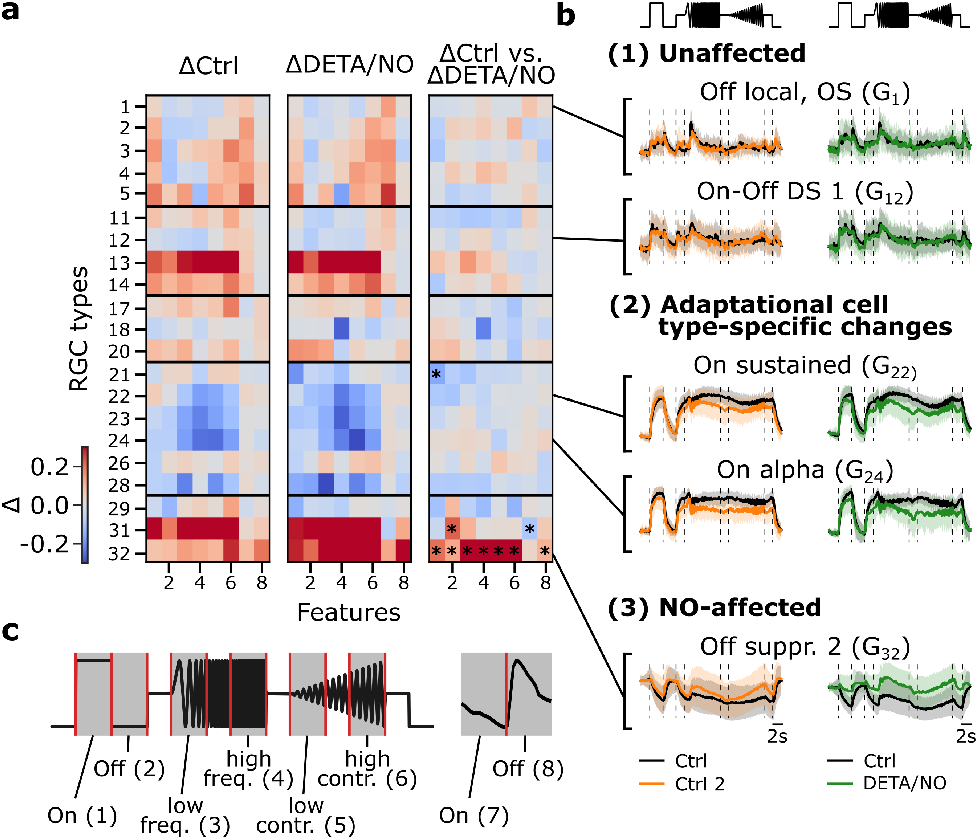
Disentangling NO-induced effects from adaptational response changes reveals type-specific NO modulation. (**a**) Left: Difference between sequentially recorded Ctrl 2 and Ctrl 1 RGC traces per type subdivided into 8 features (ΔCtrl: ΔR_Ctrl2-Ctrl1_). Color code indicates response increase (red), no change (white), and decrease (blue) for Ctrl 2. Middle: Difference between DETA/NO and Ctrl (ΔDETA/NO: ΔR_NO-Ctrl1_). Right: Difference between the two heatmaps (ΔR_NO-Ctrl1_ ΔR_Ctrl2-Ctrl1_). Asterisks indicate significant differences of the trace differences of all cells per feature and per cell type between ΔCtrl and ΔDETA/NO using independent two-sided T-test and Bonferroni correction for multiple tests; *: p<0.0003. (**b**) Example chirp traces are categorized into unaffected (top two types: G_1_, G_12_), adaptational (two middle types: G_22_, G_24_), and NO-affected (bottom: G_32_). Left traces show exemplary responses per type from the control-dataset (black: Ctrl; orange: Ctrl 2) and NO-dataset (black: Ctrl; green: DETA/NO). (**c**) Subdividing the chirp (left) and moving bar (right) stimuli into 8 features for detailed feature analysis. The chirp is subdivided into 6 features ((1) on, (2) off, (3) low frequency, (4) high frequency, (5) low contrast, and (6) high contrast); the moving bar into 2 ((7) on and (8) off).

Interestingly, a cell type from the same larger functional group that features analogous response properties, G_31_ (‘Off suppressed 1’), displayed a similar behavior during the NO application (Fig. 4a). However, G_31_ showed this response change for most features already in the control, suggesting that this effect was primarily adaptational (Fig. 4a; right panel). Still, our analysis suggests that NO had a significant effect, though different from what we observed for G_32_: in G_31_ the response to the chirp’s Off step (2) increased, whereas that to the moving bar’s leading edge (7) decreased. Note that it is possible that additional NO effects on G_31_ may have been ‘masked’ by the strong adaptational response changes.

In the previous analysis, G_18_ showed significant differences between the control- and NO-dataset, but this effect was not mirrored by the feature-wise analysis, indicating that a potential effect may be much weaker compared to the effect NO has on G_32_. Notably, RGC types that were assigned to the group of the so-called ‘Slow On’ types (G_21_G_28_), which exhibit strong and sustained responses during the frequency and contrast sequences of the chirp stimulus, showed a decrease in activity in both datasets (e.g., G_24_; Fig. 4b; (2); bottom). Consequently, the changes in these response features ((3)-(6)) are likely adaptational (Fig. 4a) — as the changes can be found in the control(Fig. 4a; left panel) as well as in the NO-dataset (Fig. 4a; middle panel), but not in the difference (Fig. 4a; right panel). Our conclusion that both adaptational and NO-mediated effects on RGC responses are highly cell type-selective is further supported by the fact that we also found several RGC types that showed stable responses during control and NO application (Fig. 4b; (1)).

Taken together, our analysis revealed that the adaptational and NO-induced effects occurred on a feature-specific as well as cell type-specific level. On the one hand, several ‘Slow On’ types showed adaptational effects in response to temporal frequency and contrast features, while other features were not affected. On the other hand, at least one distinct RGC type (G_32_) displayed a significantly increased response modulation of most features during the NO application; hence hinting towards an effect of NO on response suppression, as in the case of G_32_ is elicited by temporally changing stimulus contrast. Consequently, we focused on a more in-depth analysis of NO-induced effects on G_32_.

### Clustering of G_32_ responses reveals three functionally distinct RGC types with different NO-sensitivity

According to Baden et al. (5), G_32_ features a coverage factor of ∼4. As the average coverage factor of mouse RGCs was estimated to be ∼2 (5, 83), G_32_ likely consists of several (functional) RGC types. This is in line with the high variation of G_32_ responses, which also supports the presence of multiple functional types (see Fig. 3e).

To test this, we performed Mixture of Gaussians clustering of the RGCs assigned to G_32_ (Fig. 5a-c) using the Ctrl 1 responses to chirp and moving bar stimuli from both datasets (Fig. 5a). Since the normalized Bayesian Information Criterion (BIC; 5c, top; see Methods) values were close for n=2 and n=3 clusters, we used further tests to determine the optimal cluster number (Fig. S4a,b). These showed that for n=3 the intra-cluster correlations were higher and more consistent across clusters than for n=2 (Fig. S4c). Therefore, we concluded that G_32_ likely contains three distinct response types — all suppressed-by-contrast but to different degrees (Fig. 5d,e). All three G_32_ clusters showed little adaptation for the control-dataset, but displayed differential modulations in response to NO application, with cluster 1 exhibiting the strongest NO effect (Fig. 5f,g; left) for both stimuli. The NO effect was statistically significant in clusters 1 (n=134) and 2 (n=126) — both for mean trace difference and correlation (Fig. 5g; left and right, respectively) — but only for trace correlation in cluster 3 (n=77).

**Fig. 5.**
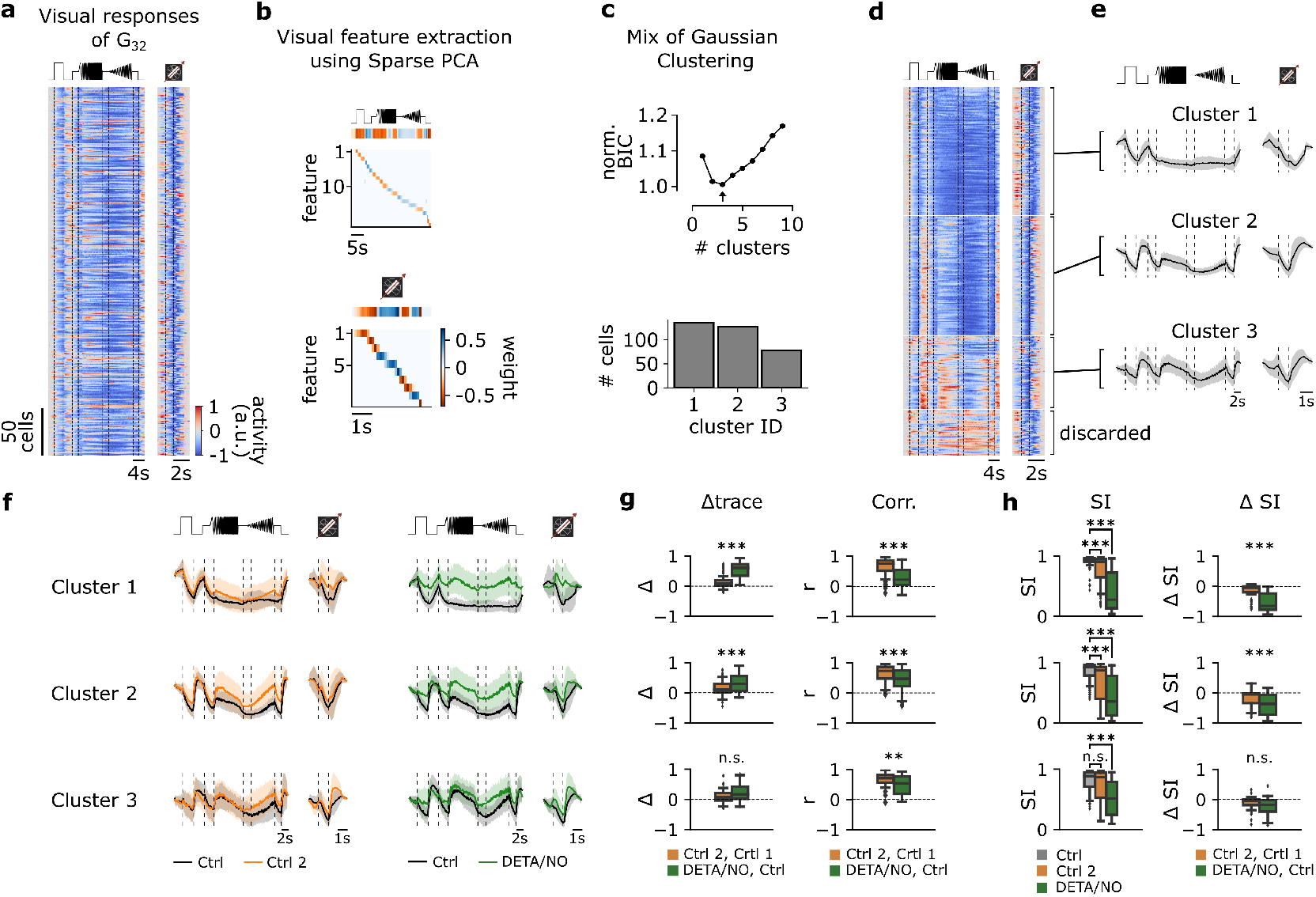
Functional clustering of the G_32_ reveals three distinct types that are differently affected by NO. (**a**) Visual responses of G_32_ cells recorded from several experiments in response to the full-field chirp (left) and moving bar (right) stimuli. (**b**) Visual features extracted from chirp (top) and moving bar (bottom) stimuli using sparse PCA on the responses. Color indicates the weight of each feature. (**c**) Top: Bayesian Information Criterion (BIC) as function of number of clusters. Arrow indicates the lowest BIC and the number of clusters to choose. Bottom: Number of cells per predicted cluster. (**d**) Cells sorted according to their assigned cluster. Cells at the bottom were discarded. (**e**) Mean responses of the 3 corresponding clusters for the chirp (left) and moving bar (right). (**f**) Left: Sequentially recorded mean responses of the 3 clusters to Ctrl 1 (black) and Ctrl 2 (orange). Right: Cluster mean responses to Ctrl 1 (black) and DETA/NO (green) (**g**) Left: Trace difference between Ctrl 2 & Ctrl 1 (orange) and DETA/NO & Ctrl (green) for the 3 clusters (clusters 1-3 from top to bottom). Right: Correlation coefficient between Ctrl 2 & Ctrl 1 (orange) and DETA/NO & Ctrl 1 (green) for the 3 clusters. n.s.: not significant; **: p<0.01, ***: p<0.001; independent T-test & Mann-Whitney U-Test. (**h**) Left: Suppression index (*SI*) computed for Ctrl 1 (gray), Ctrl 2 (orange), and DETA/NO (green) for the 3 clusters. n.s.: not significant; **: p<0.01, ***: p<0.001; Kruskal-Wallis test & Dunnett’s test. Right: Difference of *SI* between Ctrl 2 & Ctrl 1 (orange) and DETA/NO & Ctrl 1 (green). n.s.: not significant; **: p<0.01, ***: p<0.001; independent T-test & Mann-Whitney U-Test.

Since the prominent feature of these RGC types is suppression by temporal stimulus contrast, we compared their suppression strength between the conditions using a suppression index (*SI*; see Methods). Here, we found that in cluster 1, *SI* marginally, yet significantly, changed between Ctrl 1 and Ctrl 2, but was strongly and significantly reduced by NO as Δ*SI* were significantly different (Fig. 5h; top panel). Similar response behavior can be found in cluster 2, which displayed significant differences in *SI* and Δ*SI* in both control and NO-dataset (Fig. 5h; middle panel). In fact, with NO, these cells lost their suppressive feature and were rather excited than suppressed by the stimuli. In contrast, cluster 3 showed no significant differences of the *SI* between Ctrl 1 and Ctrl 2, but a clear modulation by NO (Fig. 5h; bottom panel).

Taken together, our data indicate that G_32_ may consist of three suppressed-by-contrast RGC types and that in at least two of them, the contrast suppression is strongly modulated by NO.

### NO does not affect RGC receptive field properties

So far, we focused on effects in the temporal response domain, where we found that mainly G_32_ types were affected by an increase in NO levels. However, NO has been shown to affect electrical coupling, e.g., by reducing conductance between AII ACs and On-cone BCs (66), and hence may alter receptive field (RF) properties. Therefore, we next investigated the effects of NO on the spatial RFs (sRFs) of the individual RGC types. To this end, following the same experimental paradigm as described earlier, we recorded RGC responses to binary dense noise. Next, we computed their sRFs for both recording conditions (Fig. 6a) using spike-triggered averaging (84), obtaining control and NO sRFs, and then fitted a Gaussian to each sRF’s center (Fig. 6a). We focused the following analysis on RGC types with reliable sRF estimates (see Methods). These included types in all five larger RGC groups (5): G_5_ for ‘Off’; G_11_, G_12_, and G_14_ for ‘On-Off’; G_17_ for ‘Fast On’; G_23_, G_24_, G_26_, and G_27_ for ‘Slow On’; G_31_ and G_32_ for ‘Uncertain RGCs’. In these types, sRFs were very stable in both control and NO condition.

**Fig. 6.**
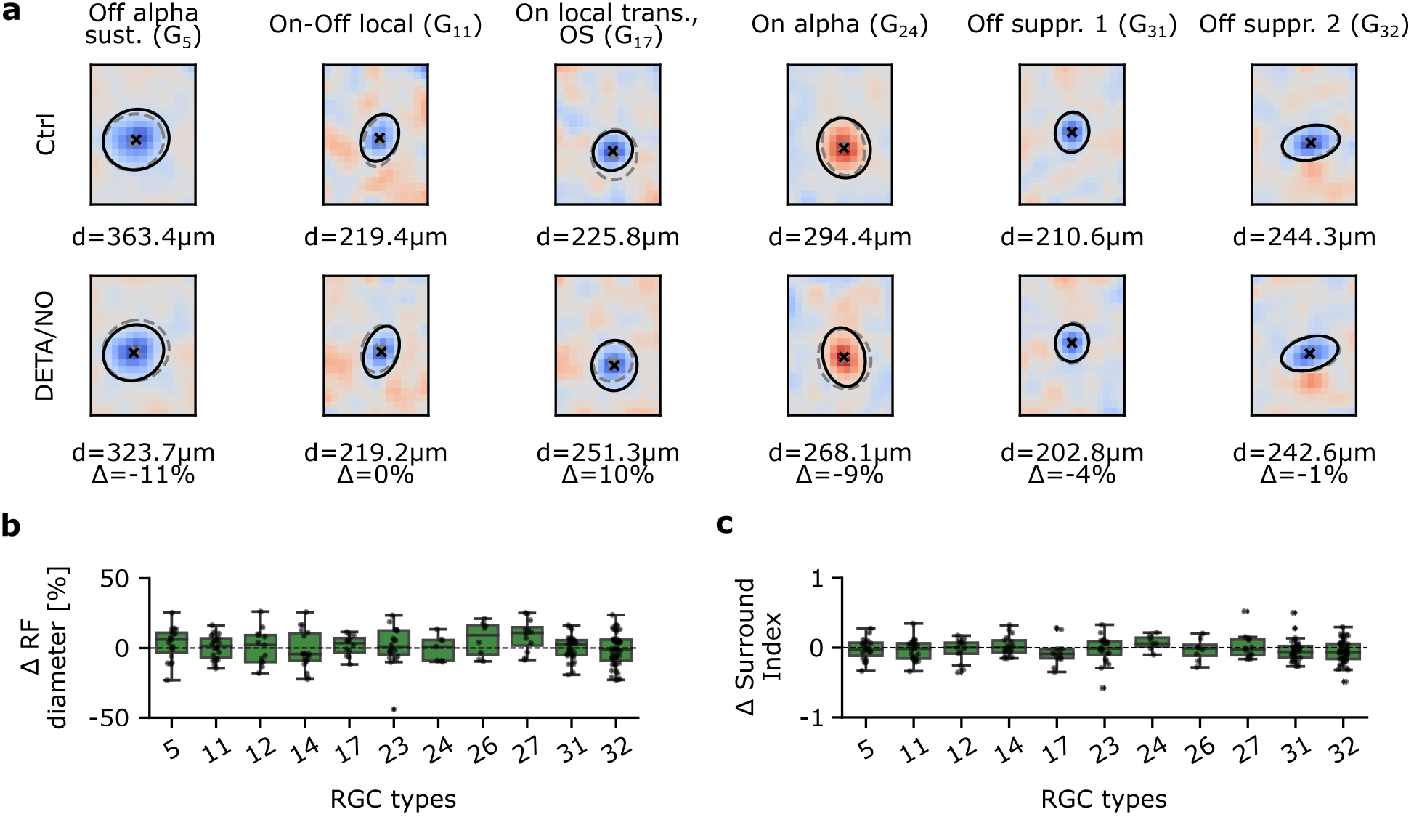
Spatial receptive fields are not affected by NO across various RGC types. (**a**) Representative estimated spatial receptive fields (sRF) of six RGC types. Top: Estimated sRFs to Ctrl 1. Cross indicates RF center; solid line indicates outline of the Gaussian fit of the RF center; dashed outline indicates corresponding Gaussian fit of the same cell to DETA/NO. Bottom: Same as top, but of DETA/NO condition. (**b**) RF diameter difference in percentage between DETA/NO and Ctrl 1. Only types with more than 5 sequentially recorded cells were included. One-sample T-test. (**c**) Surround index difference between DETA/NO and Ctrl 1. Only types with more than 5 sequentially recorded cells were included. One-sample T-test.

Using the difference in sRF center diameter between control and NO as a metric (Fig. 6b), we did not find NO to cause any significant changes in sRF size in any of the analyzed RGC types. Next, we tested if the sRF surround was affected by NO, because a modulation of inhibitory synaptic input and/or electrical coupling may cause a change in surround strength. As a measure of surround strength, we computed a surround index (see Methods) of control and NO sRFs (Fig. 6c). Like with the sRF center diameter, we did not find significant differences for any of the analyzed RGC types. Surprisingly, also G_32_ did not show NO-mediated differences in its sRF properties, implying that NO may only affect its temporal response, but not its RF organization.

Taken together, at least for the tested RGC types, we did not detect any significant NO effects neither on sRF center size nor surround strength.

### NO affects the temporal response kinetics of G_32_ subtypes

The high spatial resolution of two-photon imaging allowed us to record and identify individual RGCs. However, it is limited by having a low temporal resolution to capture subtle temporal response dynamics, which led us to record a complementary dataset using MEA recordings. Since we found that the temporal response domain of G_32_ and its subtypes were affected by NO, MEA recordings enabled us to further investigate subtle neuromodulatory effects with a higher temporal resolution.

In particular, we recorded light-induced RGC responses (n=391) from four retinae under three consecutive conditions similar to the Ca^2+^-dataset: *(1)* control, *(2)* NOdonor (DETA/NO; 100 μM) and *(3)* wash-out. As for the Ca^2+^-dataset, we used the chirp stimulus for type identification as well as a checkerboard stimulus to estimate receptive fields. To identify and compare RGC types across the Ca^2+^and MEA-dataset, we clustered the RGC responses across retinae (Fig. 7a,b) and transformed their spike trains in response to the chirp stimulus into pseudo-calcium traces as described in Goldin et al. (85) (Fig. 7c; see Methods). Next, we matched the pseudo-calcium traces with the RGC types obtained from Baden et al. (5) (Fig. 7d,e). Overall, we were able to record responses from all known RGC types, yet the sampling fractions for most types differ between the datasets, which can be explained by the recording bias inherent in MEA (86) (Fig. 7f).

**Fig. 7.**
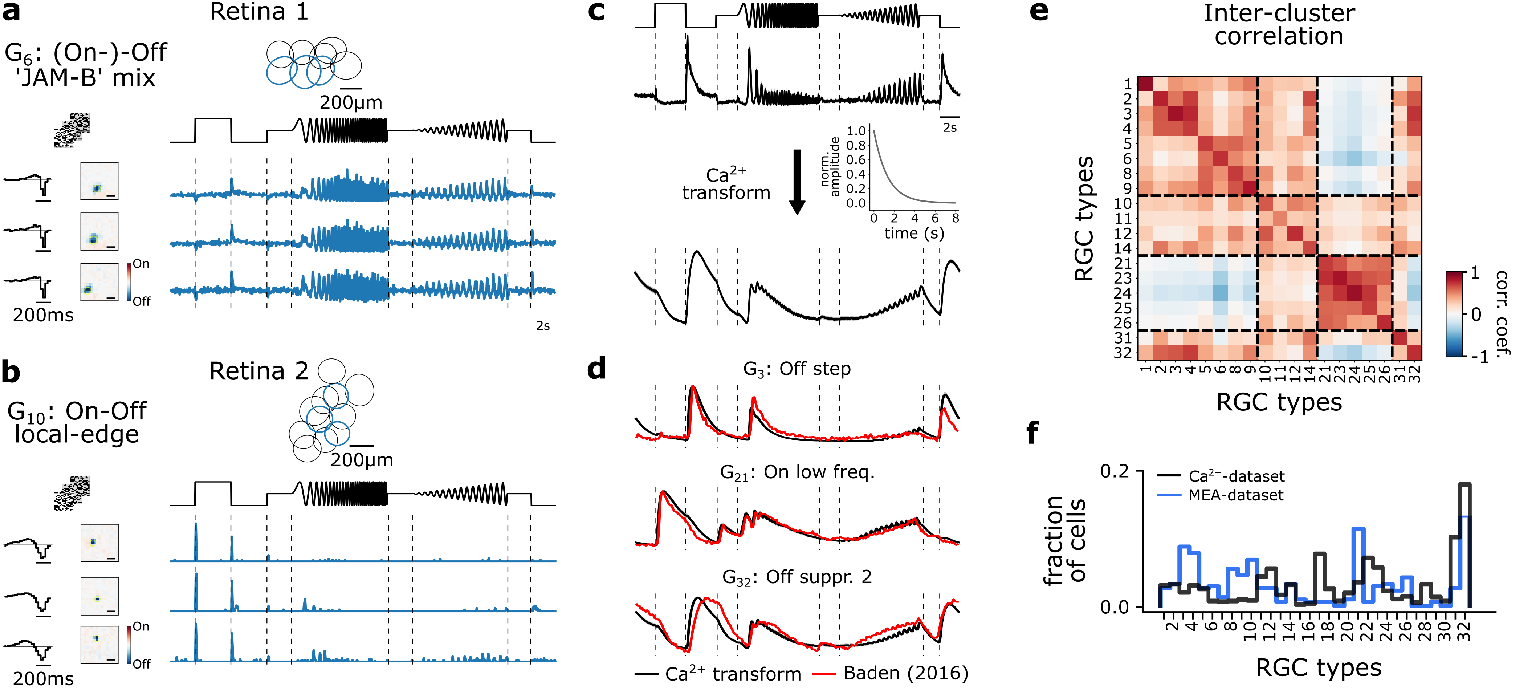
Multi-electrode recordings and pseudo-calcium transformation. (**a**) Representative RGC type, G_6_ ‘(On-)Off ‘JAM-B’ mix, recorded from one retina forming a mosaic. Three exemplary response traces of the same type. Cells are indicated in the mosaic as blue. Left: tSTAs and sSTAs computed from their responses to a checkerboard stimulus. Right: PSTHs to the chirp stimulus. (**b**) As (a)), but displaying another type, G_10_ ‘On-Off local edge’. (**c**) Illustration of the transformation of PSTHs to pseudo-calcium traces using an OGB-1 filter described in Baden et al. (5) to match and identify RGC types. (**d**) Overlay of representative mean responses of three RGC types of the Baden et al. (5) dataset (red) and the assigned Ca^2+^-transformed (black) traces to the chirp stimulus. (**e**) Inter-cluster correlation matrix of PSTHs of each assigned RGC type within a group with its group mean. Only types with n>5 cells per type were included. Dashed boxes indicate functional groups (Off, On-Off, Slow On, and Uncertain RGCs; see (5)). Color bar indicates Pearson correlation coefficient. (**f**) Comparison of the distribution of predicted RGC types between the MEA(blue) and Ca^2+^-datasets (black).

We focused the following analysis on RGCs classified as G_32_ (n=42). We found that, similar to the Ca^2+^dataset (Fig. 5), G_32_ showed reduced suppression during NO-application (Fig. 8a; top), which was not present anymore in the recording after wash-out, indicating that the NO-induced effect may be reversible (Fig. 8a; bottom). Since we found in our Ca^2+^-dataset that G_32_ consists of three subtypes, we applied the same clustering approach to the electrically recorded RGCs assigned to G_32_, revealing also three clusters (Fig. 8b, S5). For all three clusters, we found NO-induced response modulations but to varying degrees. The NO effects were at least partially reversible (see wash-out condition; Fig. 8c).

**Fig. 8.**
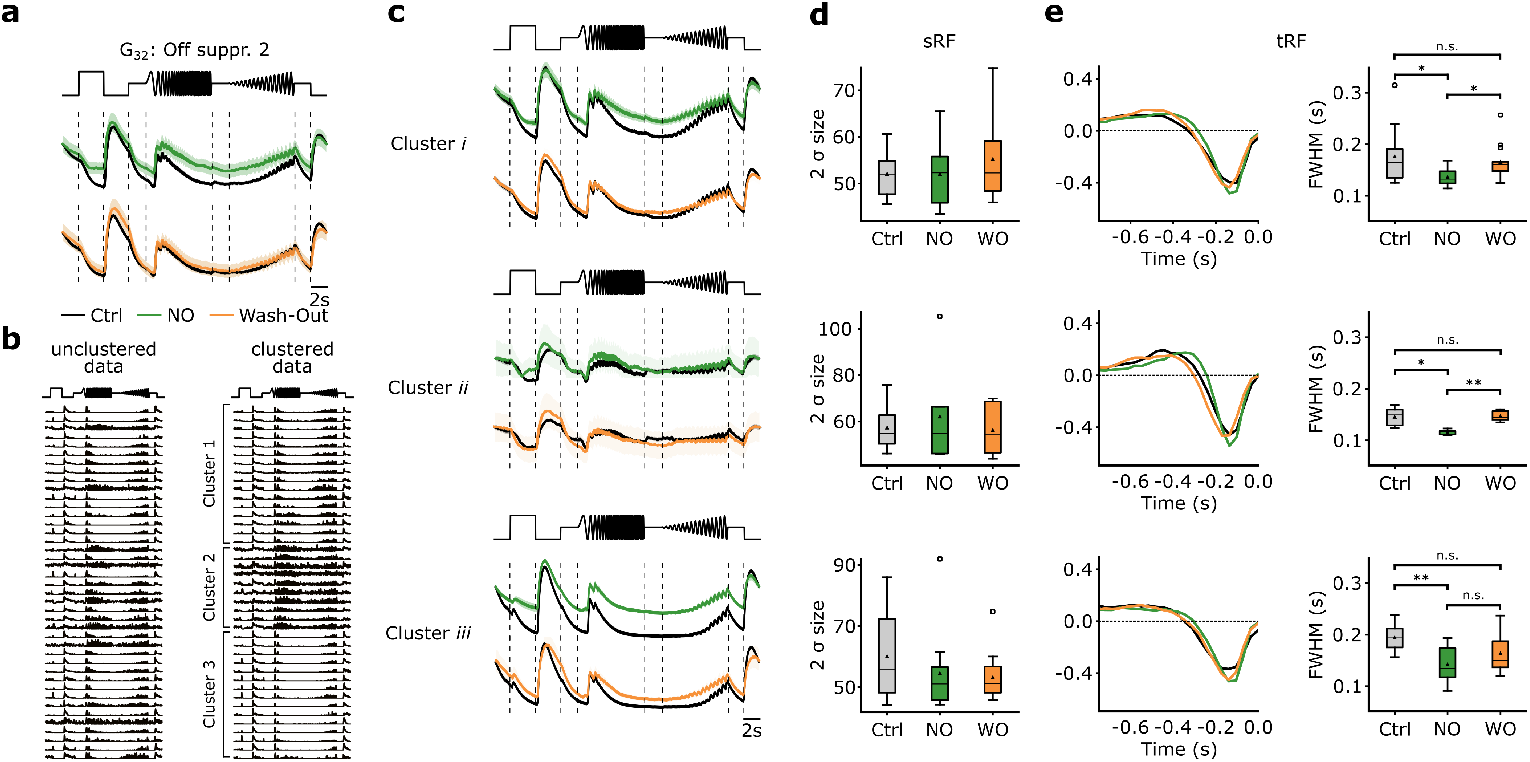
NO only affects temporal features of three distinct clusters of G_32_. (**a**) Mean Ca^2+^-transform responses to the chirp of the RGC type G_32_. Top: Sequentially recorded RGC responses to the Ctrl (black) and DETA/NO (green) conditions. Bottom: Sequentially recorded RGC responses to the Ctrl (black) and Wash-out (orange) conditions after DETA/NO application. (**b**) Left: Unclustered PSTHs of cells assigned to type G_32_. Right: Cell’s PSTHs were clustered and sorted into three distinct clusters. (**c**) Sequentially recorded mean responses of 3 clusters to Ctrl (black), DETA/NO (green), and Wash-out (orange). (**d**) Ellipse size of the fitted Gaussian of the sRF of the three conditions (Ctrl: black; DETA/NO: green; Wash-out: orange) for the 3 clusters (clusters *i* -*iii*; top to bottom). All tested conditions were not significant; Two-sided T-test. (**e**) Left: tRF kernels of the three conditions (Ctrl: black; DETA/NO: green; Wash-out: orange) for 3 clusters (clusters *i* -*iii*; top to bottom). Right: Full width at half minimum (FWHM) of the temporal RF kernels of the three conditions (Ctrl: black; DETA/NO: green; Wash-out: orange) for the 3 clusters. *: p<0.05; **: p<0.01, repeated measures ANOVA & Dunnett’s test.

We also performed an analysis directly at the spike rate level from the peristimulus time histograms (PSTH) (Fig. S6). We computed the cumulative firing rate for four time windows (features) of the chirp response, where the Ca^2+^-dataset suggests suppression (Fig. S6b,c). We found for all three clusters an increase in cumulative firing rate during DETA/NO application — indicative of a reduction in suppression — for at least one feature (Fig. S6c). That the NO effect was differential (e.g., compare ‘Freq. step’ and ‘On Step’ between clusters) further supports the presence of multiple RGC types in G_32_.

Based on the Ca^2+^-dataset, we did not detect significant effects of NO on RFs. However, as recording time, and hence stimulus presentation time, is limited per scan field, we repeated the RF analysis for the MEA-dataset. MEA recordings allow for much longer dense noise stimuli and, thus, can yield more precise RF estimates. In line with the Ca^2+^-dataset, we did not find any NO-induced modulation of sRF size in any of the clusters (Fig. 8d). However, the temporal RF (tRF) kernels showed significantly faster response kinetics during NO-application for all three clusters (Fig. 8e). Also, the effects were reversible for two clusters (except for cluster *iii*). That we did not detect this effect in the Ca^2+^-dataset is likely due to the lower temporal resolution of the Ca^2+^ recordings. Note that the G_32_ sub-types identified in both datasets do not necessarily correspond to the same RGC types: While two clusters show high correlations between Ca^2+^ and MEA data (clusters 2 vs. *ii*, 3 vs. *i*), one clearly differs in its temporal response (cluster 1 vs. *iii*; Fig S7a,b; see also Methods).

Taken together, our MEA-dataset confirms that G_32_ consists of three sub-types and that their temporal responses are modulated by NO: it reduced their suppression by temporal contrast. Additionally, the analysis of the MEA-dataset (tRF) revealed that all three sub-types displayed faster response kinetics under NO, an effect that was reversible. The sRFs were unchanged by NO, in line with the Ca^2+^ data.

## Discussion

We used two-photon Ca^2+^ imaging and MEA recordings to measure RGC responses to various visual stimuli to investigate the neuromodulatory effects of elevated NO levels on signal processing across RGC types in the mouse retina. To our surprise, even without pharmacological perturbation, we found that about one-third of the RGC types displayed highly reproducible and cell type-specific response changes during the course of an experiment – a finding of potentially high relevance especially for pharmacological experiments in the *ex vivo* retina using two-photon Ca^2+^ imaging. Accounting for these adaptational changes enabled us to isolate NO-related effects on RGC responses. Here, we revealed that mainly the RGCs assigned to G_32_ (‘Off suppressed 2’) were affected by NO, which strongly reduced the response suppression and rendered the cells more active. Further, we demonstrated that G_32_ likely consists of three types — consistent with its high coverage factor (5) — that were all differentially modulated by NO. Finally, for a representative subset of RGC types, we showed that elevating the NO-level had no discernible effect on sRF size or surround strength. In addition, we were able to confirm those results using MEA recordings and showed that NO caused faster temporal response kinetics of these three subtypes and partially increased firing rates; these changes were mostly reversible. Together, our data suggest that NO specifically modulates response suppression and kinetics in a group of contrast-suppressed RGC types. Additionally, our study demonstrates the need for recording paradigms that take adaptational, non-drug-related response changes into account when analyzing potentially subtle pharmacological effects.

## Nitric oxide as a (neuro-)modulator in the retina

Neuronal nitric oxide synthesis (nNOS) has been detected in different retinal cell classes (49–57) and the main NO-sensor (NO-GC), which connects NO to intracellular cGMP signaling (55, 60, 87, 88), is present in all retinal layers (70). This and earlier findings of NO modulating the response gain of BCs (68, 69) and RGCs (71) suggest that NO constitutes a neuromodulatory system within the retina involved in light adaptation. However, in the light-adapted mouse retina, light stimulus-dependent NO production seems to mainly occur in specific AC types, with the nNOS-2 AC being the main source of endogenous NO (36). In particular, it releases NO in response to flickering light (i.e. fast changes in contrast) (24, 36, 55, 60, 72). Thus, it has been proposed that nNOS-2 ACs report fast contrast changes at photopic conditions (24, 36). This points at an interesting though speculative functional role of the NO-sensitivity in G_32_, namely that NO helps “highlighting” *changes* in contrast by reducing suppression and accelerating response kinetics. Thereby, G_32_ RGCs could relay contrast changes to higher visual targets.

Interestingly, this RGC type-selectivity of NO is reminiscent of a recent study, where the neuromodulator dopamine was found to modulate distinct response features of specific RGC subtypes (48). What cellular mechanisms underlie the action of NO on G_32_ RGCs, or its upstream circuit, will be interesting to investigate in future studies.

To ensure reliable RGC type classification, our set of visual stimuli was restricted to artificial ones (full-field chirps, moving bars, and dense noise). Such artificial stimuli probe the stimulus space in a rather selective and limited fashion. Hence, we cannot exclude that we missed NO effects that may have become apparent for other, more complex stimuli. Specifically, natural images or movies — stimuli that are closer to what the retina evolved to process (89) — may be needed for a more complete picture of the functional implication of retinal neuromodulation.

That natural stimuli can reveal novel nonlinear properties of retinal functions was demonstrated, for example, by Goldin et al. (85), who showed that an RGC’s contrast selectivity can be context-dependent. Similarly, Höfling et al. (90) recorded natural movie-evoked RGC responses to train a convolutional neural network, which through *in silico* experiments allowed them to discover that transient suppressed-by-contrast RGC (G_28_) feature center color-opponent responses and may signal context changes. These studies highlight that future studies on retinal neuromodulators should also employ natural stimuli.

Another important aspect to consider when studying NO neuromodulation in the retina is the level of light adaptation. Several studies proposed that NO facilitates the transition across light levels (66, 68, 69), especially to photopic conditions (36). Since we employed two-photon imaging, which inevitably results in a certain level of background ‘illumination’ (see discussion in (76, 91)), our experiments were performed in the low photopic range. Therefore, NO-mediated neuromodulation may serve additional light-level dependent functions: More globally during the transition from scotopic to mesopic/photopic, and more cell typespecific in the photopic regime, as we reported here.

Finally, a gaseous neuromodulator like NO poses a additional problem when studying its effects: A donor is needed and the final concentration of the neuromodulator delivered to the cells depends on many factors (92). We used the NO-donor DETA/NO, because its long half-life time (*t*_1*/*2_) of > 20 hrs enables a steady delivery of NO within a tissue. Assuming for NO in tissue a *t*_1*/*2_ of 2 min, a freshly prepared DETA/NO solution of 100 μM is expected to release about 0.25 μM NO (93). The endogenous NO concentration in the retina was estimated to be a few 100 nM (e.g., (94, 95), which means that the NO concentration we applied is likely within the physiological range.

It has been reported for DETA/NO that the donor itself — independent of NO — may affect the electrical properties of cultured cerebellar granule cells by reversibly activating a cation-selective channels (96). While we cannot exclude that this side-effect of DETA/NO may have contributed to the effects we observed in G_32_ RGCs, we consider this unlikely, because substantial side-effects were mainly observed by Thompson et al. (96) at much higher DETA/NO concentrations (3 mM) than we used in our study.

### Suppressed-by-Contrast cells in the mouse retina

Functionally, RGC types referred to as “Suppressed-by-Contrast” (SbC) are characterized by a decrease in their activity for both positive and negative temporal contrasts within their RF (97–100). In Baden et al. (5), three functional RGC types were labeled SbC based on their light stimulus-evoked Ca^2+^ response being suppressed primarily by positive (On-SbC: G_28_) or negative temporal contrast (Off-SbCs: G_31_, G_32_).

Notably, G_32_ (“Off suppressed 2”) is also suppressed by the moving bar stimulus, suggesting that the cells are also sensitive to spatial contrast (i.e. an edge appearing in their RF). Coverage analysis indicated that G_32_ may contain several RGC types (5) — in line with our cluster analysis. It is still unclear if G_32_ contains one (or more) of the individually studied SbC types in mouse (97, 98, 100, 101), yet, recently, Goetz et al. (6) speculated if the novel bursty-SbC RGC (101) aligns with (a sub-cluster of) G_32_.

### Adaptational, cell type-specific response changes

Every recording method introduces technique-specific biases that have to be considered in the data analysis and interpretation. For two-photon imaging with fluorescent Ca^2+^ sensors, these potential biases include Ca^2+^ buffering, sensor bleaching by the excitation laser, and, in the case of bulk-loading with synthetic dyes (as in our experiments; see also (102)), slow leakage of the indicator from the cells. In retinal imaging, additional potentially confounding factors are an excitation laser-induced baseline activity and photoreceptor bleaching (77, 91). These biases are expected to be systematic, e.g., causing a general decrease in signal-to-noise ratio across (RGC) responses. To account for this, our recording paradigm produced a controland a NO-dataset consisting of sequentially recorded RGC responses.

When analyzing the control-dataset, we were surprised by finding response changes in the 2^nd^ control measurement 10 min after the 1^st^ control measurement in approx. one third of the RGC types in the absence of the NO-donor. These changes were consistent for a particular type but differed between types. Notably, we did not observe a simple overall decrease or increase in activity, but rather selective changes of response features: For instance, in G_24_ (‘Slow On’) only the response to high frequency and low contrast was reduced, while the remaining response features were not affected. As mentioned above, the time differences between Ctrl 1 & Ctrl 2 was 10 min and Ctrl & DETA/NO was approx. 25 min. However, we do not think that this has a substantial effect on our results because when a change for either Ctrl 2 or DETA/NO was observed, it followed the same trend — other than in G_32_ (and G_18_).

Together, this strongly argues against a systematic, recording technique-related bias but rather for an adaptational effect. Currently, we can only speculate about the mechanism(s) underlying this type-specific adaptation. The most parsimonious explanation may be related to the *ex vivo* condition of the retina: While we allowed the tissue to settle and adapt to perfusion medium, light level, temperature, etc. for ∼25 min, extracellular signaling molecules, such as neuromodulators, may be depleted and washed out throughout the experiment, resulting in differential adaptation of various RGC types. In any case, as type-selective adaptations can confound the recorded responses in a complex manner, a sequential-recording paradigm as the one described here, is recommended — in particular for pharmacological experiments.

### Combining large-scale population recordings, RGC classification, and sequential recordings to study neuromodulation of retinal output

In this study, we investigated the neuromodulatory effects of NO on the retinal output signal. To this end, we combined experimental and computational approaches to dissect NO-mediated effects at the RGC type-level. The latter is important for understanding neuromodulator function for early vision because the visual information is sent to the brain via parallel feature channels, represented by *>*40 RGC types in the mouse retina (5, 6, 83, 103). We demonstrated that our approach enabled us to distinguish adaptational from actual NO-induced effects using a rather simple and straightforward linear analysis (i.e., focusing on mean trace or RF size differences), assuming that the adaptational- and NO-effects are independent and sum linearly, which means that we may have missed potential nonlinear effects.

The combination of two-photon Ca^2+^ imaging and MEA recordings allowed us to confirm our findings and make use of two methods that complement one another. Finally, the presented pipeline for analyzing neuromodulation in neural circuits constitutes a framework that can be easily extended by applying more advanced analyses.

## ACKNOWLEDGEMENTS

We thank Gordon Eske & Merle Harrer for technical support and Clint L. Makino, Philipp Berens & Robert Feil for fruitful discussions. This work was funded by the Deutsche Forschungsgemeinschaft (DFG, German Research Foundation) 335549539/GRK2381 and CRC 1233 (project number 276693517). M. Goldin was funded by a fellowship from Fondation de France and by CNRS and by grant ANR PerBaCo. We acknowledge support from the Open Access Publication Fund of the University of Tübingen.

## Methods

### Animals and tissue preparation for two-photon Ca^2+^ imaging

All animals for the two-photon Ca^2+^ imaging experiments were conducted at the University of Tübingen and were performed according to the laws governing animal experimentation issued by the German Government as well as approved by the institutional animal welfare committee of the University of Tübingen. For all experiments, we used retinae (n=26) from C57Bl/6J mice (n=14; JAX 000664) of either sex between the age of 4-16 weeks. All animals were kept in the local animal facility and housed under the standard 12h/12h day/night cycle at 22°C and a humidity of 55%.

The following procedures were carried out under very dim red (> 650 nm) light. Before each imaging experiment, the animal was dark-adapted for >1h, then anesthetized with isoflurane (CP-Pharma) and sacrificed by cervical dislocation. Immediately after, the eyes were enucleated with a dorsal cut as orientation landmark and hemisected in carboxygenated (95% O_2_, 5% CO_2_) artificial cerebrospinal fluid (ACSF) solution containing (in mM): 125 NaCl, 2.5 KCl, 2 CaCl_2_, 1 MgCl_2_, 1.25 NaH_2_PO_4_, 26 NaHCO_3_, 20 glucose, and 0.5 L-glutamine at pH 7.4. Sulforhodamine-101 (SR101, 0.1 μM; Invitrogen) was added to the ACSF to reveal blood vessels and damaged ganglion cell layer (GCL) cells in the red fluorescence channel (76). The carboxygenated ACSF was constantly perfused through the recording chamber at 4 ml/min and kept at ∼36°C throughout the entire experiment. After the dissection, retinae were bulk-electroporated with the synthetic fluorescent calcium indicator Oregon-Green 488 BAPTA-1 (OGB-1; hexapotassium salt; Life Technologies) (102). To electroporate the GCL, the dissected retina was flat-mounted with the GCL facing up onto an AnodiscTM (#13, 0.1 μm pore size, 13 mm diameter, Cytiva), and then placed between two 4 mm horizontal platinum disk electrodes (CUY700P4E/L, Nepagene/Xceltis). The lower electrode was covered with 15 μl of ACSF, while a 10 μl drop of 5 mM OGB-1 dissolved in ACSF was suspended from the upper electrode and lowered onto the retina. Then, nine electrical pulses (∼9.2 V, 100 ms pulse width, at 1 Hz) from a pulse generator/wideband amplifier combination (TGP110 and WA301, Thurlby handar/Farnell) were applied and then, the electroporated retina on the Anodisc was transferred into the recording chamber, whereby the dorsal edge of the retina pointed away from the experimenter. The retina was left there for ∼30 min to recover, as well as adapted to the light stimulation by displaying a binary dense noise stimulus (20 × 15 matrix, 40 × 40 μm^2^ pixels, balanced random sequence) at 5 Hz before the recordings started.

### Two-photon Ca^2+^ imaging

For the functional Ca^2+^ imaging experiments, a MOM-type two-photon microscope (designed by W. Denk, MPI, Heidelberg; purchased from Sutter Instruments/Science Products) (76, 77) was employed. The microscope was equipped with a mode-locked Ti:Sapphire laser (MaiTai-HP DeepSee, Newport SpectraPhysics) tuned to 927 nm (ideal wavelength to excite OGB1), two photomultiplier tubes serving as fluorescence detection channels for OGB-1 (HQ 510/84, AHF/Chroma) and SR101 (HQ 630/60, AHF), and a water immersion objective (CF175 LWD×16/0.8W, DIC N2, Nikon, Germany). To acquire images, custom-made software (ScanM by M. Müller and T. Euler) running under IGOR Pro 6.3 for the operating system Microsoft Windows (Wavemetrics) was used and time-lapsed 64 × 64 pixel image scans (100 × 100 μm) at 7.8125 Hz were taken. Routinely, the optic nerve position and the scan field position were recorded to reconstruct their retinal positions. High-resolution images (512 × 512 pixel images) were recorded to support semi-automatic ROI detection.

### Light stimulation for two-photon Ca^2+^ imaging

For the light stimulation of the retinal tissue, a digital light processing (DLP) projector (lightcrafter (LCr), DPME4500UVBGMKII, EKB Technologies Ltd) was used to display the visual stimuli through the objective onto the retina, whereby the stimulus was focused on the photoreceptor layer (104). The LCr was equipped with a lightguide port to couple in external, band-pass filtered green and UV light-emitting diode (LEDs; green: 576 BP 10, F37-576; UV: 387 BP 11, F39-387; both AHF/Chroma). The band-pass filter was used to optimize the spectral separation of mouse M- and Sopsins (390/576 Dualband, F59-003, AHF/Chroma). Both LEDs were synchronized with the scan retracing of the microscope. Stimulator intensity (as photoisomerization rate, 10^3^ P^*\**^s^*−*1^ per cone) was calibrated to range from ∼0.5 (black image) to ∼20 for M-and S-opsins, respectively. Steady illumination of ∼10^4^ P^*\**^s^*−*1^ per cone was present during the scan recordings due to the two-photon excitation of photopigments (76, 77).

In total, three types of light stimuli were used for the imaging of Ca^2+^ in the GCL: (1) full-field chirp stimulus (700 μm ∅ see details here ref. (5)), (2) bright moving bar (0.3 × 1 mm) at 1 mm s^*−*1^ in eight directions to probe direction and orientation selectivity, and (3) random binary noise with a checkerboard grid of 20 × 15 checks and a check size of 40 μm at 5 Hz for 5 min to map receptive fields. Light stimulus center and scan field center were aligned. Before each stimulus was presented, the baseline was recorded after the laser started scanning for at least 30 s to avoid immediate laser-induced effects on the retinal activity (76, 77, 105).

### Animals and tissue preparation for electrophysiological recordings

Electrophysiological data were recorded from isolated retinae from four C57Bl/6J mice of 8 - 10 weeks. The experiment was performed in accordance with the institutional animal care standards of Sorbonne Université. (Paris, France). The animals were housed in enriched cages with ad libitum food, and watering. The ambient temperature was between 22 and 25°C, the humidity was between 50 and 70%, and the light cycle was 12–14h of light, 10–12h of darkness. After killing the animal, the eye was enucleated and transferred rapidly into oxygenated Ames medium (Sigma-Aldrich). Dissection was made under dim light condition as described previously (106, 107).

### Electrophysiological recordings

For the electrophysiological recordings, we mounted a piece of retina onto a membrane and then lowered it with the ganglion cell side against a 252-channel multi-electrode array (MEA) whose electrodes were spaced by 30 μm. During dissection and recordings, the tissue was perfused with oxygenated Ames solution and a peristaltic perfusion system with two independent pumps: PPS2 (Multichannel Systems GmbH). Mice retinae were kept at 35–37°C during the whole experiment. The data sampling rate was 20 kHz. The raw signal was acquired through MC Rack Multi-channel Systems software 4.6.2, it was high-pass filtered at 100 Hz, and the spikes were isolated using SpyKING CIRCUS 1.0.628 (106). Subsequent data analysis was done with customwritten Python codes. We extracted the activity of a total of 391 neurons. We kept cells with a low number of refractory period violations (< 0.95%, with median < 0.05% for all experiments, 2 ms refractory period) and whose template waveform could be well distinguished from the template waveforms of other cells. These constraints ensured a good quality of the reconstructed spike trains. In addition, we discarded neurons that showed no or almost no responses to chirp stimulus, preventing the correct cell classification.

### Light stimulation for electrophysiological recordings

A white-mounted LED (MCWHLP1, Thorlabs Inc.) was used as a light source, and the stimuli were displayed using a Digital Mirror Device (DLP9500, Texas Instruments) and focused on the photoreceptors using standard optics and an inverted microscope (Nikon). The light level corresponded to photopic vision: 4.9 × 10^4^ and 1.4 × 10^5^ isomerizations / (photoreceptor.s) for S-cones and M-cones, respectively. In total, we displayed two stimuli: *(1)* a random binary checkerboard with check size of 42 μm for 30-40 min at 30 Hz, and *(2)* a full field chirp stimulus as used for the twophoton Ca^2+^ imaging experiments. It was played at 50 Hz, containing 20 repetitions of 32 s length.

### Pharmacology

All used drugs were added to the carboxygenated, perfused ACSF solution 15 min prior to the second recording of the GCL scan fields. For the drugs, the respective concentrations were used (in μM): 100 (Z)-1-[N-(2-aminoethyl)-N-(2-ammonioethyl)amino]diazen-1-ium-1,2-diolate (DETA/NO) and 1 strychnine. The ACSF solution with and without drug application was always kept at ∼36°C.

### Data Analysis of two-photon Ca^2+^ imaging recordings

Image extraction and semi-automatic region-of-interest (ROI) detection were performed using Igor PRO 8. All analyses were organized and performed in a custom-written schema using DataJoint for Python http://datajoint.github.io/; D. Yatsenko, Tolias lab, Baylor College of Medicine.

#### Preprocessing

After the Ca^2+^ were extracted from individual ROIs, as described elsewhere (5, 105). The raw traces were detrended by subtracting a smoothed version *r*_*smooth*_ of the trace from the raw one. Detrending was necessary to remove slow drifts in the signal that were unrelated to the light-induced response. The smoothed trace *r*_*smooth*_ was computed by applying a Savitzky-Golay filter (108) of 3^*rd*^ polynomial order and a window length of 60 s using the Python SciPy implementation *scipy*.*signal*.*savgol*_*f ilter* (109).

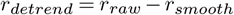

For the chirp and moving bar stimulus, detrended traces were averaged over repetitions; and in the case of the moving bar stimulus, reduced to the response average for the preferred motion direction of the cell (for details see Baden et al. (5)). Finally, response averages were normalised by first subtracting the baseline activity (computed as the mean over the first second), and then by dividing by the maximum amplitude *max*_*t*_(|*r*(*t*)|) = 1. This normalization was performed independently for each ROI, stimulus, and condition.

#### Inclusion criterion

To include reliable cell responses for the performed analyses, two consecutive quality filtering steps were applied. At first, the response quality criterion, also termed quality index (QI), was computed for the moving bar (QI_MB_ > 0.6) and full-field chirp (QI_chirp_ > 0.45). Cells that passed either one of these two QIs in both recording conditions were included, otherwise they were discarded in the following analyses. As in Baden et al. (5), the quality index is defined as follows:

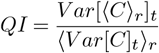

where *C* is the *T* by *R* response matrix (time samples by stimulus repetitions) and ⟨⟩_*x*_ and *V ar*[]_*x*_ denote the mean and variance across the indicated dimension *x*, respectively. As a second step, cells were assigned to an RGC group using the RGC classifier, which returned the RGC group index and a confidence score (i.e., assignment probability to the predicted RGC group by the random forest classifier (80)). Only cells that were assigned to one of the RGC groups (i.e., RGC index 1 - 32) were included, whereas cells assigned to a displaced AC group (i.e., RGC index 33 - 46) were rejected. Cells that exceeded the confidence score threshold of > 0.25 were included.

#### Suppression index

For each cell, the suppression index (SI) was measured by comparing the (absolute) negative area under the curve (AUC_*neg*_) of the chirp and moving bar responses with the total area under the curve (AUC_*neg*_+AUC_*pos*_) of the entire response trace. For the absolute AUC_*neg*_, the response was clipped for value <0.

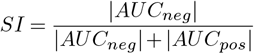

#### Trace difference Δ

To compute the trace difference between the sequentially recorded responses per cell, we subtracted the first response average to the chirp and moving bar stimuli, i.e., Ctrl 1, from the second recorded light-induced response, i.e., either NO or Ctrl 2. For the cell type-specific analysis, we computed the average trace differences per cell. For the feature-based analysis, we computed the average trace differences per feature and per cell type.

#### On-Off Index

The On-Off index was computed as

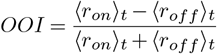

where *r*_*on*_ and *r*_*off*_ are defined as the separated time components of the moving bar response into its On- and Off-component. For each component, we computed the mean value of the discrete differences along the time axis clipped between 0 and 1 to estimate if there is a response to the particular feature.

#### Classification of functional retinal ganglion cell types

For the functional classification of RGC types, we used a previously published RGC classifier (80). The classifier, which uses a random forest classifier, was trained, validated, and tested on previously published RGC type responses (5). As input to the classifier, we used the responses to the standard set of stimuli, i.e., full-field chirp and moving bar, as well as soma sizes (separates alpha and non-alpha types) and the p-values of the permutation test for direction selectivity (separates DS and Non-DS types). For every cell, the RGC clas-sifier outputs its type index and the confidence scores for all 46 types. The confidence score, as described in ‘inclusion criterion’ was used as a quality criterion.

#### Receptive field estimation

We mapped receptive fields (RFs) of RGCs using the RF python toolbox *RFEst* (110), following the procedure in (5) with few modifications. The binary dense noise stimulus (20 × 15 matrix, (40 μm)^2^ pixels, balanced random sequence; 5 Hz) was centered on the recording field. We computed the temporal gradients of the Ca^2+^ signals from the detrended traces and clipped negative values:

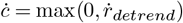

The stimulus *S*(*t*) and the clipped temporal gradients *c* were upsampled to 10 times the stimulus frequency to compute the gradient-triggered average stimulus:

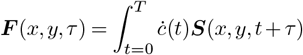

where ***S***(*x, y, t*) is the stimulus, *τ* is the lag ranging from approximately −0.20 to 1.38 seconds, and *T* is the duration of the stimulus. We smoothed these raw RF using a 5 × 5 pixel and 1 pixel standard deviation Gaussian window for each lag. Then we decomposed the RF into a temporal (*F*_*t*_(*τ*)) and spatial (***F***_*s*_(*x, y*)) component using singular value decomposition and scaled them such that max(|*F*_*t*_|) = 1 and max(|***F***_*s*_|) = max(|***F*** |). RF quality was computed as:

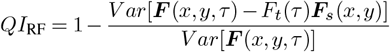

Only RFs with *QI*_RF_ *>* 0.45 were used for the analysis. For each spatial RF ***F***_*s*_, we fit a 2D Difference of Gaussians using the python package *astropy* (111). The mean and covariance matrices of the center and surround Gaussian fits were tied, except for a linear scaling of the covariance matrix. We defined the polarity *p* ∈ {−1, 1} of the spatial RF as the sign of the model fit at its mean. Next, we computed the center RF of the spatial RF as 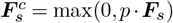. The surround index was computed as:

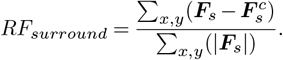

To measure the center RF size, we fit a 2D Gaussian to the center RF, with the mean fixed to the one obtained from the Difference of Gaussians fit. The area covered by two standard deviations of this Gaussian fit was used as the RF size.

#### Functional clustering

The functional clustering was based on a similar approach as in (5). The clustering was only applied on RGC types previously classified as G_32_ and only recorded in Ctrl 1. First, visual features from the full-field chirp and moving bar Ca^2+^ responses were extracted using sparse principal component analysis (112). After optimizing the alpha parameter for each stimulus, each cell’s dimensionality was reduced to 30 features, whereby the chirp covered 20 and the moving bar 10 features. Alpha was optimized in a way that every part of each stimulus was represented by one feature to increase interpretability. Each feature was standardized across cells before clustering. Then, the features were used to cluster the cells using a Mixture of Gaussian model. The ideal number of clusters was chosen based on the cross-validated Bayesian Information Criterion. Additionally, cluster coherence was computed and validated using intraand inter-cluster correlation, as well as the influence of potential batch effects, i.e., a single cluster originates from a single retina or scan field, but is found across several ones. Then, the model was used to predict cell type labels. Finally, cluster 3 showed a high signal-to-noise ratio, thus cells were re-clustered, which originated in three clusters, whereby two showed high variability in their chirp and moving bar responses. These cells were discarded in the further analysis to clean this cluster from potential contamination.

#### Statistical analysis

To quantify the differences between traces, a Shaprio-Wilk test was used to test for normality and then either the two-sided t-test (if normally distributed) or the non-parametric Wilcoxon signed-rank test (Mann-Whitney U Test). To determine *α*, Bonferroni correction was used, depending on the number of tests performed. To test the difference between traces against zero, we either used the t-test or the non-parametric Wilcoxon signed-rank test, depending on the distribution. A one-sampled T-test was performed to test the mean against a population mean of zero to quantify if the mean difference diverges from zero. For the statistical comparison of the suppression index between conditions per cluster, the non-parametric KruskalWallis one-way analysis of variance and post-hoc Dunnett’s test and Bonferroni correction to determine the statistical significance between conditions were used.

### Data Analysis of electrophysiological recordings

#### Functional clustering

To cluster cells in different functional types, we based our analysis on the chirp and checkerboard stimulus responses and represented each RGC with a reduced vector. To obtain these vectors, first, we constructed peri-stimulus histograms (PSTH) from the spikes evoked from the chirp stimulus, using a binning of 100 ms, and spike-triggered averages (STA). Then, for each experiment, we z-scored all PSTHs and performed a principle component analysis (PCA) on them. We kept the number of components that were needed to explain 80% variance of the data (11-12 components for the PSTH). Second, we used the temporal profile (40 samples at 30 Hz) of each cell’s STA obtained using the checkerboard stimulus. We z-scored it and performed a PCA, keeping two components, which explains around 80% of the variance. This adds information about the classical STA polarity of the RGCs. Third, we used the area of the ellipse fitted to the classical STA, as the product value of their major and minor axis σ values. These areas were normalized from 0 to 1. In this way, we obtain a data vector of around 15 values, depending on the experiment, that describes each RGC according to their response to a chirp and a checkerboard. Then, we performed an agglomerative clustering, setting the threshold value in a way that all clusters looked homogeneous across PSTHs and STAs. This resulted in over-clustering producing between 12 and 44 RGC groups (from 29 to 149 cells in each experiment).

#### Receptive field estimation

To estimate spatial and temporal receptive fields, we displayed a random binary checkerboard with check size of 42 μm for 30-40 min at 30 Hz. A three-dimensional spike-triggered average (STA; x, y, and time) was sampled using 40 time samples. The spatial STA presented across all the figures was obtained as the 2-dimensional spatial slice at the maximum value after smoothing. The temporal STA is the one-dimensional time slice at that same value. A double Gaussian fit was performed on the resulting spatial STA, and the ellipse corresponding to a 2σ contour of the fit was plotted for all the figures.

#### Pseudo-calcium transformation

In the last step, we assigned each cluster to one of the 32 types described in Baden et al. (5). To do this, we used the Ca^2+^-dataset to match it with the MEA-dataset. We based our analysis on their reported data in the Extended data from Fig. 1, where the authors link electrophysiology and Ca^2+^ imaging by means of a convolution between a Ca^2+^ event-triggered by a single spike.

We transformed the PSTHs by convolving them with a decaying exponential (see Fig. 7c; (85)), in which we adjusted the temporal decay constant to maximize the correlation of our clusters and theirs (median maximum correlation of 0.78). RGC types that presented strong responses to the modulating frequencies and amplitude were assigned correctly, while other types, which mostly respond to ON/OFF steps, were assigned in a second round of correlation match after excluding the former clusters. Besides the correlation of the chirp traces, we confirmed the correct assigning of clusters by checking that the ellipses of each type form a proper mosaic (see Fig. 7a,b), that the spatial STAs look uniform, and the similarity of their spike waveforms. After doing this for every experiment, we grouped the cells of each type across animals where we found a clear homogeneity. Finally, we computed a correlation matrix between the average chirp response of each type to show that the clusters are homogeneous (see Fig. 7e).

With this procedure, we were able to match a majority of RGC types between those datasets, yet aligning the datasets is challenging. In fact, the cell types underlying the Ca^2+^ and MEA RGC clusters may not always be same. A caveat is that while Ca^2+^ is a proxy for spiking activity, other Ca^2+^ sources as well as sub-threshold membrane potential changes may affect the intracellular Ca^2+^, potentially in a cell type-specific way.

#### Functional cell typing of G_32_

Once we obtained all the cells assigned to G_32_, we pooled the data from the 4 experiments and performed the same steps as before, to further distinguish sub-groups as we did with the Ca^2+^ imaging data. We first over-clustered again in this step and merged the similar sub-clusters in a last step. We obtained again, now in the electrophysiological data, three clearly differentiated subgroups. From an initial set of 46 cells, we obtained three subgroups of 17, 10, and 15 cells (Fig. 8c), plus a nonhomogeneous subgroup of four cells that did not fit into any of them and was discarded.

#### Statistical analysis

To quantify the ellipses of the sRFs, we used the two-sides T-test. To quantify if the FWHM values of the tRFs were significantly different, we performed repeated measures ANOVA and post-hoc Dunnett’s test. Only cells where we could compute sRFs or tRFs across the three conditions were used for these analyses.

## Data and code availability

Data, light stimuli, as well as all custom analyses including code and notebooks to reproduce analyses will be made available at https://github.com/eulerlab upon journal publication.

## Author contributions

D.G., Z.Z., T.E., and T.S. designed the study; T.S. wrote the animal protocol with the help of D.G. and T.E.; D.G. performed the imaging and electrophysiological experiments, supported by M.A.G., R.A., and T.S.K.; D.G. performed pre-processing; D.G. analyzed the data with the help of J.O. and M.A.G.; Z.Z. helped to maintain the experimental setup; D.G. wrote the manuscript with the help of T.E., T.S., J.O., M.A.G., and O.M.

## Conflicts of interest

The authors declare no conflicts of interest.

## Notes

### Competing Interest Statement

The authors have declared no competing interest.

### Summary of Updates

We modified parts of the results, added several supplementary figures and revised the discussion.

## Bibliography

1. Heinz Wässle and Brian B Boycott. Functional architecture of the mammalian retina. Physiological reviews, 71(2):447–480, 1991.

2. Richard H Masland. The fundamental plan of the retina. Nature neuroscience, 4(9): 877–886, 2001.

3. Heinz Wässle. Parallel processing in the mammalian retina. Nature Reviews Neuroscience, 5(10):747–757, 2004.

4. Tania A Seabrook, Timothy J Burbridge, Michael C Crair, and Andrew D Huberman. Architecture, function, and assembly of the mouse visual system. Annual review of neuroscience, 40:499–538, 2017.

5. Tom Baden, Philipp Berens, Katrin Franke, Miroslav Román Rosón, Matthias Bethge, and Thomas Euler. The functional diversity of retinal ganglion cells in the mouse. Nature, 529:345–350, 2016. ISSN 9781137332875. doi: 10.1038/nature16468.

6. Jillian Goetz, Zachary F. Jessen, Anne Jacobi, Adam Mani, Sam Cooler, Davon Greer, Sabah Kadri, Jeremy Segal, Karthik Shekhar, Joshua R. Sanes, and Gregory W. Schwartz. Unified classification of mouse retinal ganglion cells using function, morphology, and gene expression. Cell Rep, 40(2):111040, 2022. ISSN 2211-1247 (Electronic). doi: 10.1016/j.celrep.2022.111040.

7. James R Anderson, Bryan W Jones, Carl B Watt, Margaret V Shaw, Jia-Hui Yang, David DeMill, James S Lauritzen, Yanhua Lin, Kevin D Rapp, David Mastronarde, et al. Exploring the retinal connectome. Molecular vision, 17:355, 2011.

8. Robert E Marc, Bryan W Jones, Carl B Watt, James R Anderson, Crystal Sigulinsky, and Scott Lauritzen. Retinal connectomics: towards complete, accurate networks. Progress in retinal and eye research, 37:141–162, 2013.

9. Moritz Helmstaedter, Kevin L Briggman, Srinivas C Turaga, Viren Jain, H Sebastian Seung, and Winfried Denk. Connectomic reconstruction of the inner plexiform layer in the mouse retina. Nature, 500(7461):168–174, 2013.

10. Moritz Helmstaedter. Cellular-resolution connectomics: challenges of dense neural circuit reconstruction. Nature methods, 10(6):501–507, 2013.

11. Felice A Dunn and Rachel OL Wong. Wiring patterns in the mouse retina: collecting evidence across the connectome, physiology and light microscopy. The Journal of physiology, 592(22):4809–4823, 2014.

12. Jamie Johnston and Leon Lagnado. General features of the retinal connectome determine the computation of motion anticipation. Elife, 4:e06250, 2015.

13. Kevin L Briggman, Moritz Helmstaedter, and Winfried Denk. Wiring specificity in the direction-selectivity circuit of the retina. Nature, 471(7337):183–188, 2011.

14. Huayu Ding, Robert G Smith, Alon Poleg-Polsky, Jeffrey S Diamond, and Kevin L Briggman. Species-specific wiring for direction selectivity in the mammalian retina. Nature, 535(7610):105–110, 2016.

15. Paul Witkovsky. Dopamine and retinal function. Documenta ophthalmologica, 108:17– 39, 2004.

16. Suva Roy and Greg D Field. Dopaminergic modulation of retinal processing from starlight to sunlight. Journal of pharmacological sciences, 140(1):86–93, 2019.

17. Rebekah A Warwick, Serena Riccitelli, Alina S Heukamp, Hadar Yaakov, Lea Ankri, Jonathan Mayzel, Noa Gilead, Reut Parness-Yossifon, and Michal Rivlin-Etzion. Top-down modulation of the retinal code via histaminergic neurons of the hypothalamus. bioRxiv, pages 2022–04, 2022.

18. Ursula Greferath, M Kambourakis, Christian Barth, E. Fletcher, and M Murphy. Characterization of histamine projections and their potential cellular targets in the mouse retina. Neuroscience, 158(2):932–944, 2009.

19. Justine Masson. Serotonin in retina. Biochimie, 161:51–55, 2019.

20. Stephen Yazulla. Endocannabinoids in the retina: from marijuana to neuroprotection. Progress in retinal and eye research, 27(5):501–526, 2008.

21. Thomas Schwitzer, Raymund Schwan, Karine Angioi-Duprez, Anne Giersch, and Vincent Laprevote. The endocannabinoid system in the retina: from physiology to practical and therapeutic applications. Neural plasticity, 2016, 2016.

22. Charles F Yates, Jin Y Huang, and Dario A Protti. Tonic endocannabinoid levels modulate retinal signaling. International Journal of Environmental Research and Public Health, 19 (19):12460, 2022.

23. Ira M Goldstein, Philipp Ostwald, and Steven Roth. Nitric oxide: a review of its role in retinal function and disease. Vision research, 36(18):2979–2994, 1996.

24. Alex H Vielma, Mauricio A Retamal, and Oliver Schmachtenberg. Nitric oxide signaling in the retina: what have we learned in two decades? Brain research, 1430:112–125, 2012.

25. Ana Santos-Carvalho, Ana Rita Alvaro, Joao Martins, Antonio Francisco Ambrosio, and Claudia Cavadas. Emerging novel roles of neuropeptide y in the retina: from neuromodulation to neuroprotection. Progress in neurobiology, 112:70–79, 2014.

26. Ana Santos-Carvalho, António Francisco Ambrósio, and Cláudia Cavadas. Neuropeptide y system in the retina: From localization to function. Progress in retinal and eye research, 47:19–37, 2015.

27. Wenjun Yan, Mallory A Laboulaye, Nicholas M Tran, Irene E Whitney, Inbal Benhar, and Joshua R Sanes. Mouse retinal cell atlas: molecular identification of over sixty amacrine cell types. Journal of Neuroscience, 40(27):5177–5195, 2020.

28. Matthew J Gastinger, Ning Tian, Tamas Horvath, and David W Marshak. Retinopetal axons in mammals: emphasis on histamine and serotonin. Current eye research, 31 (7-8):655–667, 2006.

29. Donald M O’Malley and Richard H Masland. Co-release of acetylcholine and gamma-aminobutyric acid by a retinal neuron. Proceedings of the National Academy of Sciences, 86(9):3414–3418, 1989.

30. Donald M O’Malley, Julie H Sandell, and Richard H Masland. Co-release of acetylcholine and gaba by the starburst amacrine cells. Journal of Neuroscience, 12(4):1394–1408, 1992.

31. Hajime Hirasawa, Rebecca A Betensky, and Elio Raviola. Corelease of dopamine and gaba by a retinal dopaminergic neuron. Journal of Neuroscience, 32(38):13281–13291, 2012.

32. Jeffrey S Diamond. Inhibitory interneurons in the retina: types, circuitry, and function. Annual review of vision science, 3:1–24, 2017.

33. Michael D Flood, Johnnie M Moore-Dotson, and Erika D Eggers. Dopamine d1 receptor activation contributes to light-adapted changes in retinal inhibition to rod bipolar cells. Journal of neurophysiology, 120(2):867–879, 2018.

34. Manvi Goel and Stuart C Mangel. Dopamine-mediated circadian and light/dark-adaptive modulation of chemical and electrical synapses in the outer retina. Frontiers in Cellular Neuroscience, 15:647541, 2021.

35. Rolf Herrmann, Stephanie J Heflin, Timothy Hammond, Bowa Lee, Jing Wang, Raul R Gainetdinov, Marc G Caron, Erika D Eggers, Laura J Frishman, Maureen A McCall, et al. Rod vision is controlled by dopamine-dependent sensitization of rod bipolar cells by gaba. Neuron, 72(1):101–110, 2011.

36. Jason Jacoby, Amurta Nath, Zachary F Jessen, and Gregory W Schwartz. A self-regulating gap junction network of amacrine cells controls nitric oxide release in the retina. Neuron, 100(5):1149–1162, 2018.

37. B Bauer, B Ehinger, and L Åberg. [3 h]-dopamine release from the rabbit retina. Albrecht von Graefes Archiv für klinische und experimentelle Ophthalmologie, 215:71–78, 1980.

38. Bernard F Godley and Richard J Wurtman. Release of endogenous dopamine from the superfused rabbit retina in vitro: effect of light stimulation. Brain research, 452(1-2): 393–395, 1988.

39. Mustafa B. A. Djamgoz and Hans-Joachim Wagner. Localization and function of dopamine in the adult vertebrate retina. Neurochemistry international, 20(2):139–191, 1992.

40. Víctor Pérez-Fernández, Nina Milosavljevic, Annette E Allen, Kirstan A Vessey, Andrew I Jobling, Erica L Fletcher, Paul P Breen, John W Morley, and Morven A Cameron. Rod photoreceptor activation alone defines the release of dopamine in the retina. Current Biology, 29(5):763–774, 2019.

41. Charles W Nichols, David Jacobowitz, and Marianne Hottenstein. The influence of light and dark on the catecholamine content of the retina and choroid. Investigative Ophthalmology & Visual Science, 6(6):642–646, 1967.

42. Izhak Nir, Rashidul Haque, and P Michael Iuvone. Diurnal metabolism of dopamine in the mouse retina. Brain research, 870(1-2):118–125, 2000.

43. Christophe Ribelayga, Yu Cao, and Stuart C Mangel. The circadian clock in the retina controls rod-cone coupling. Neuron, 59(5):790–801, 2008.

44. W Wade Kothmann, Stephen C Massey, and John O’Brien. Dopamine-stimulated dephosphorylation of connexin 36 mediates aii amacrine cell uncoupling. Journal of Neuroscience, 29(47):14903–14911, 2009.

45. Nan Ge Jin and Christophe P Ribelayga. Direct evidence for daily plasticity of electrical coupling between rod photoreceptors in the mammalian retina. Journal of Neuroscience, 36(1):178–184, 2016.

46. Eric M Lasater and John E Dowling. Dopamine decreases conductance of the electrical junctions between cultured retinal horizontal cells. Proceedings of the National Academy of Sciences, 82(9):3025–3029, 1985.

47. Yuki Hayashida, Carolina Varela Rodríguez, Genki Ogata, Gloria J Partida, Hanako Oi, Tyler W Stradleigh, Sherwin C Lee, Anselmo Felipe Colado, and Andrew T Ishida. Inhibition of adult rat retinal ganglion cells by d1-type dopamine receptor activation. Journal of Neuroscience, 29(47):15001–15016, 2009.

48. Rebekah A Warwick, Alina S Heukamp, Serena Riccitelli, and Michal Rivlin-Etzion. Dopamine differentially affects retinal circuits to shape the retinal code. The Journal of physiology, 2023.

49. David S Bredt, Paul M Hwang, and Solomon H Snyder. Localization of nitric oxide synthase indicating a neural role for nitric oxide. Nature, 347(6295):768–770, 1990.

50. Ted M Dawson, David S Bredt, Majid Fotuhi, Paul M Hwang, and Solomon H Snyder. Nitric oxide synthase and neuronal nadph diaphorase are identical in brain and peripheral tissues. Proceedings of the National Academy of Sciences, 88(17):7797–7801, 1991.

51. Ryo Yamamoto, David S. Bredt, Solomon H. Snyder, and Richard A. Stone. The localization of nitric oxide synthase in the rat eye and related cranial ganglia. Neuroscience, 54(1):189–200, 1993.

52. Ryo Yamamoto, David S Bredt, Ted M Dawson, Solomon H Snyder, and Richard A Stone. Enhanced expression of nitric oxide synthase by rat retina following pterygopalatine parasympathetic denervation. Brain research, 631(1):83–88, 1993.

53. Todd A Blute, Bernd Mayer, and William D Eldred. Immunocytochemical and histochemical localization of nitric oxide synthase in the turtle retina. Visual Neuroscience, 14(4): 717–729, 1997.

54. Todd A Blute, Michael R Lee, and William D Eldred. Direct imaging of nmda-stimulated nitric oxide production in the retina. Visual Neuroscience, 17(4):557–566, 2000.

55. William D Eldred and Todd A Blute. Imaging of nitric oxide in the retina. Vision research, 45(28):3469–3486, 2005.

56. Vikram Palamalai, Ruth M Darrow, Daniel T Organisciak, and Masaru Miyagi. Light-induced changes in protein nitration in photoreceptor rod outer segments. Mol Vis, 12: 1543–1551, 2006.

57. Jan J Blom, Todd A Blute, and William D Eldred. Functional localization of the nitric oxide/cgmp pathway in the salamander retina. Visual Neuroscience, 26(3):275–286, 2009.

58. Silke Haverkamp and Heinz Wässle. Immunocytochemical analysis of the mouse retina. Journal of Comparative Neurology, 424(1):1–23, 2000.

59. Thomas J Giove, Monika M Deshpande, and William D Eldred. Identification of alternate transcripts of neuronal nitric oxide synthase in the mouse retina. Journal of neuroscience research, 87(14):3134–3142, 2009.

60. Jan J Blom, Tom Giove, Monika Deshpande, and William D Eldred. Characterization of nitric oxide signaling pathways in the mouse retina. Journal of Comparative Neurology, 520(18):4204–4217, 2012.

61. Gerard P Ahern, Vitaly A Klyachko, and Meyer B Jackson. cgmp and s-nitrosylation: two routes for modulation of neuronal excitability by no. Trends in neurosciences, 25(10): 510–517, 2002.

62. Ari Sitaramayya. Soluble guanylate cyclases in the retina. Molecular and cellular biochemistry, 230:177–186, 2002.

63. John Garthwaite. Dynamics of cellular no-cgmp signaling. Frontiers in Bioscience-Landmark, 10(2):1868–1880, 2005.

64. Masaru Miyagi, Hirokazu Sakaguchi, Ruth M Darrow, Lin Yan, Karen A West, Kulwant S Aulak, Dennis J Stuehr, Joe G Hollyfield, Daniel T Organisciak, and John W Crabb. Evidence that light modulates protein nitration in rat retina. Molecular & Cellular Proteomics, 1(4):293–303, 2002.

65. Douglas T Hess, Akio Matsumoto, Sung-Oog Kim, Harvey E Marshall, and Jonathan S Stamler. Protein s-nitrosylation: purview and parameters. Nature reviews Molecular cell biology, 6(2):150–166, 2005.

66. Stephen L Mills and Stephen C Massey. Differential properties of two gap junctional pathways made by aii amacrine cells. Nature, 377(6551):734–737, 1995.

67. Scott Nawy and Craig E Jahr. Suppression by glutamate of cgmp-activated conductance in retinal bipolar cells. Nature, 346(6281):269–271, 1990.

68. Josefin Snellman and Scott Nawy. cgmp-dependent kinase regulates response sensitivity of the mouse on bipolar cell. Journal of Neuroscience, 24(29):6621–6628, 2004.

69. Ryan E Tooker, Mikhail Y Lipin, Valerie Leuranguer, Eva Rozsa, Jayne R Bramley, Jacqueline L Harding, Melissa M Reynolds, and Jozsef Vigh. Nitric oxide mediates activity-dependent plasticity of retinal bipolar cell output via s-nitrosylation. Journal of Neuroscience, 33(49):19176–19193, 2013.

70. Alex H Vielma, Adolfo Agurto, Joaquín Valdés, Adrián G Palacios, and Oliver Schmachtenberg. Nitric oxide modulates the temporal properties of the glutamate response in type 4 off bipolar cells. PLoS One, 9(12):e114330, 2014.

71. Guo-Yong Wang, Lauren C Liets, and Leo M Chalupa. Nitric oxide differentially modulates on and off responses of retinal ganglion cells. Journal of neurophysiology, 90(2): 1304–1313, 2003.

72. Guo-Yong Wang, Deborah A Van der List, Joseph P Nemargut, Julie L Coombs, and Leo M Chalupa. The sensitivity of light-evoked responses of retinal ganglion cells is decreased in nitric oxide synthase gene knockout mice. Journal of Vision, 7(14):7–7, 2007.

73. Joseph P Nemargut and Guo-Yong Wang. Inhibition of nitric oxide synthase desensitizes retinal ganglion cells to light by diminishing their excitatory synaptic currents under light adaptation. Vision research, 49(24):2936–2947, 2009.

74. Iqbal Ahmad, Trese Leinders-Zufall, Jeffery D Kocsis, Gordon M Shepherd, Frank Zufall, and Colin J Barnstable. Retinal ganglion cells express a cgmp-gated cation conductance activatable by nitric oxide donors. Neuron, 12(1):155–165, 1994.

75. Fusao Kawai and Peter Sterling. cgmp modulates spike responses of retinal ganglion cells via a cgmp-gated current. Visual neuroscience, 19(3):373–380, 2002.

76. Thomas Euler, Susanne E. Hausselt, David J. Margolis, Tobias Breuninger, Xavier Castell, Peter B. Detwiler, and Winfried Denk. Eyecup scope—optical recordings of light stimulus-evoked fluorescence signals in the retina. Pflügers Archiv - European Journal of Physiology, 457:1393–1414, 2009. doi: 10.1007/s00424-008-0603-5.

77. Thomas Euler, Katrin Franke, and Tom Baden. Studying a light sensor with light: Multiphoton imaging in the retina. e. hartveit (ed.). Multiphoton Microscopy, Neuromethods (Humana, New York, NY), 148:225–250, 2019. doi: 10.1007/978-1-4939-9702-2_10.

78. Zhijian Zhao, David A Klindt, André Maia Chagas, Klaudia P Szatko, Luke Rogerson, Dario A Protti, Christian Behrens, Deniz Dalkara, Timm Schubert, Matthias Bethge, et al. The temporal structure of the inner retina at a single glance. Scientific reports, 10(1):1– 17, 2020.

79. Dominic Gonschorek, Larissa Höfling, Klaudia P Szatko, Katrin Franke, Timm Schubert, Benjamin Dunn, Philipp Berens, David Klindt, and Thomas Euler. Removing inter-experimental variability from functional data in systems neuroscience. Advances in Neural Information Processing Systems, 34:3706–3719, 2021.

80. Yongrong Qiu, David A. Klindt, Klaudia P. Szatko, Dominic Gonschorek, Larissa Hoefling, Timm Schubert, Laura Busse, Matthias Bethge, and Thomas Euler. Efficient coding of natural scenes improves neural system identification. bioRxiv, page 2022.01.10.475663, 2022. doi: 10.1101/2022.01.10.475663.

81. Jan Weiss, Gregory A. O’sullivan, Liane Heinze, H-X Chen, Heinrich Betz, and Heinz Wässle. Glycinergic input of small-field amacrine cells in the retinas of wildtype and glycine receptor deficient mice. Molecular and Cellular Neuroscience, 37(1):40–55, 2008.

82. Reza Farajian, Feng Pan, Abram Akopian, Béla Völgyi, and Stewart A Bloomfield. Masked excitatory crosstalk between the on and off visual pathways in the mammalian retina. The Journal of physiology, 589(18):4473–4489, 2011.

83. J Alexander Bae, Shang Mu, Jinseop S Kim, Nicholas L Turner, Ignacio Tartavull, Nico Kemnitz, Chris S Jordan, Alex D Norton, William M Silversmith, Rachel Prentki, et al. Digital museum of retinal ganglion cells with dense anatomy and physiology. Cell, 173 (5):1293–1306, 2018.

84. EJ Chichilnisky. A simple white noise analysis of neuronal light responses. Network: computation in neural systems, 12(2):199, 2001.

85. Matías A Goldin, Baptiste Lefebvre, Samuele Virgili, Mathieu Kim Pham Van Cang, Alexander Ecker, Thierry Mora, Ulisse Ferrari, and Olivier Marre. Context-dependent selectivity to natural images in the retina. Nature Communications, 13(1):5556, 2022.

86. Francesco Trapani, Giulia Lia Beatrice Spampinato, Pierre Yger, and Olivier Marre. Differences in nonlinearities determine retinal cell types. Journal of Neurophysiology, 130 (3):706–718, 2023.

87. Todd A Blute, Paula Velasco, and William D Eldred. Functional localization of soluble guanylate cyclase in turtle retina: modulation of cgmp by nitric oxide donors. Visual neuroscience, 15(3):485–498, 1998.

88. Sebastian Gotzes, Jan de Vente, and Frank Mueller. Nitric oxide modulates cgmp levels in neurons of the inner and outer retina in opposite ways. Visual neuroscience, 15(5): 945–955, 1998.

89. Yongrong Qiu, Zhijian Zhao, David Klindt, Magdalena Kautzky, Klaudia P Szatko, Frank Schaeffel, Katharina Rifai, Katrin Franke, Laura Busse, and Thomas Euler. Natural environment statistics in the upper and lower visual field are reflected in mouse retinal specializations. Current Biology, 31(15):3233–3247, 2021.

90. Larissa Höfling, Klaudia P Szatko, Christian Behrens, Yuyao Deng, Yongrong Qiu, David Alexander Klindt, Zachary Jessen, Gregory W Schwartz, Matthias Bethge, Philipp Berens, et al. A chromatic feature detector in the retina signals visual context changes. eLife, 13:e86860, 2024.

91. Thomas Euler, Susanne E Hausselt, David J Margolis, Tobias Breuninger, Xavier Castell, Peter B Detwiler, and Winfried Denk. Eyecup scope—optical recordings of light stimulus-evoked fluorescence signals in the retina. Pflügers Archiv-European Journal of Physiology, 457:1393–1414, 2009.

92. Joseph S Beckman and Willem H Koppenol. Nitric oxide, superoxide, and peroxynitrite: the good, the bad, and ugly. American Journal of Physiology-cell physiology, 271(5): C1424–C1437, 1996.

93. Anand Ramamurthi and Randy S Lewis. Measurement and modeling of nitric oxide release rates for nitric oxide donors. Chemical research in toxicology, 10(4):408–413, 1997.

94. Micah Guthrie. Measurement of intraretinal nitric oxide in early diabetic retinopathy. Illinois Institute of Technology, 2014.

95. GR Kalamkarov, AE Bugrova, TS Konstantinova, and TF Shevchenko. Endogenous nitric oxide content in the cellular layers of the retina. Neuroscience and Behavioral Physiology, 46:291–295, 2016.

96. AJ Thompson, PK Mander, and GC Brown. The no donor deta-nonoate reversibly activates an inward current in neurones and is not mediated by the released nitric oxide. British journal of pharmacology, 158(5):1338–1343, 2009.

97. Nai-Wen Tien, James T Pearson, Charles R Heller, Jay Demas, and Daniel Kerschensteiner. Genetically identified suppressed-by-contrast retinal ganglion cells reliably signal self-generated visual stimuli. Journal of Neuroscience, 35(30):10815–10820, 2015.

98. Nai-Wen Tien, Tahnbee Kim, and Daniel Kerschensteiner. Target-specific glycinergic transmission from vglut3-expressing amacrine cells shapes suppressive contrast responses in the retina. Cell reports, 15(7):1369–1375, 2016.

99. Jason Jacoby, Yongling Zhu, Steven H DeVries, and Gregory W Schwartz. An amacrine cell circuit for signaling steady illumination in the retina. Cell reports, 13(12):2663–2670, 2015.

100. Jason Jacoby and Gregory William Schwartz. Typology and circuitry of suppressed-by-contrast retinal ganglion cells. Frontiers in Cellular Neuroscience, 12:269, 2018.

101. Sophia Wienbar and Gregory William Schwartz. Differences in spike generation instead of synaptic inputs determine the feature selectivity of two retinal cell types. Neuron, 110 (13):2110–2123, 2022.

102. Kevin L Briggman and Thomas Euler. Bulk electroporation and population calcium imaging in the adult mammalian retina. Journal of neurophysiology, 105(5):2601–2609, 2011.

103. Nicholas M Tran, Karthik Shekhar, Irene E Whitney, Anne Jacobi, Inbal Benhar, Guosong Hong, Wenjun Yan, Xian Adiconis, McKinzie E Arnold, Jung Min Lee, et al. Single-cell profiles of retinal ganglion cells differing in resilience to injury reveal neuroprotective genes. Neuron, 104(6):1039–1055, 2019.

104. Katrin Franke, André Maia Chagas, Zhijian Zhao, Maxime JY Zimmermann, Philipp Bartel, Yongrong Qiu, Klaudia P Szatko, Tom Baden, and Thomas Euler. An arbitrary-spectrum spatial visual stimulator for vision research. elife, 8:e48779, 2019.

105. Klaudia P Szatko, Maria M Korympidou, Yanli Ran, Philipp Berens, Deniz Dalkara, Timm Schubert, Thomas Euler, and Katrin Franke. Neural circuits in the mouse retina support color vision in the upper visual field. Nature communications, 11(1):3481, 2020.

106. Pierre Yger, Giulia LB Spampinato, Elric Esposito, Baptiste Lefebvre, Stéphane Deny, Christophe Gardella, Marcel Stimberg, Florian Jetter, Guenther Zeck, Serge Picaud, et al. A spike sorting toolbox for up to thousands of electrodes validated with ground truth recordings in vitro and in vivo. Elife, 7:e34518, 2018.

107. Olivier Marre, Dario Amodei, Nikhil Deshmukh, Kolia Sadeghi, Frederick Soo, Timothy E Holy, and Michael J Berry. Mapping a complete neural population in the retina. Journal of Neuroscience, 32(43):14859–14873, 2012.

108. William H Press and Saul A Teukolsky. Savitzky-golay smoothing filters. Computers in Physics, 4(6):669–672, 1990.

109. Pauli Virtanen, Ralf Gommers, Travis E. Oliphant, Matt Haberland, Tyler Reddy, David Cournapeau, Evgeni Burovski, Pearu Peterson, Warren Weckesser, Jonathan Bright, Stéfan J. van der Walt, Matthew Brett, Joshua Wilson, K. Jarrod Millman, Nikolay Mayorov, Andrew R. J. Nelson, Eric Jones, Robert Kern, Eric Larson, C J Carey, İlhan Polat, Yu Feng, Eric W. Moore, Jake VanderPlas, Denis Laxalde, Josef Perktold, Robert Cim-rman, Ian Henriksen, E. A. Quintero, Charles R. Harris, Anne M. Archibald, Antônio H. Ribeiro, Fabian Pedregosa, Paul van Mulbregt, and SciPy 1.0 Contributors. SciPy 1.0: Fundamental Algorithms for Scientific Computing in Python. Nature Methods, 17:261– 272, 2020. doi: 10.1038/s41592-019-0686-2.

110. Ziwei Huang, Yanli Ran, Jonathan Oesterle, Thomas Euler, and Philipp Berens. Estimating smooth and sparse neural receptive fields with a flexible spline basis. arXiv preprint arXiv:2108.07537, 2021.

111. Adrian M Price-Whelan, BM Sipo?cz, H. Günther, PL Lim, SM Crawford, S Conseil, DL Shupe, MW Craig, N Dencheva, A Ginsburg, et al. The astropy project: Building an open-science project and status of the v2. 0 core package. The Astronomical Journal, 156(3):123, 2018.

112. Hui Zou, Trevor Hastie, and Robert Tibshirani. Sparse principal component analysis. Journal of computational and graphical statistics, 15(2):265–286, 2006.

