## Supplementary Figures for "Nitric oxide modulates contrast suppression in a subset of mouse retinal ganglion cells"

1448 # Supplementary Material

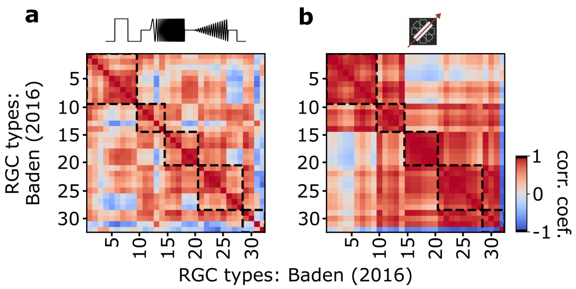


**Fig. S1. Autocorrelation matrix of Baden et al. (2016) dataset. (a)** Autocorrelation matrix of average type responses per RGC types of the Baden et al. (2016) dataset for responses to the chirp stimulus. Dashed boxes indicate functional groups (Off, On-Off, Fast On, Slow On, and Uncertain RGCs). Color bar indicates Pearson correlation coefficient. **b** As (a), but in response to the moving bar stimulus.

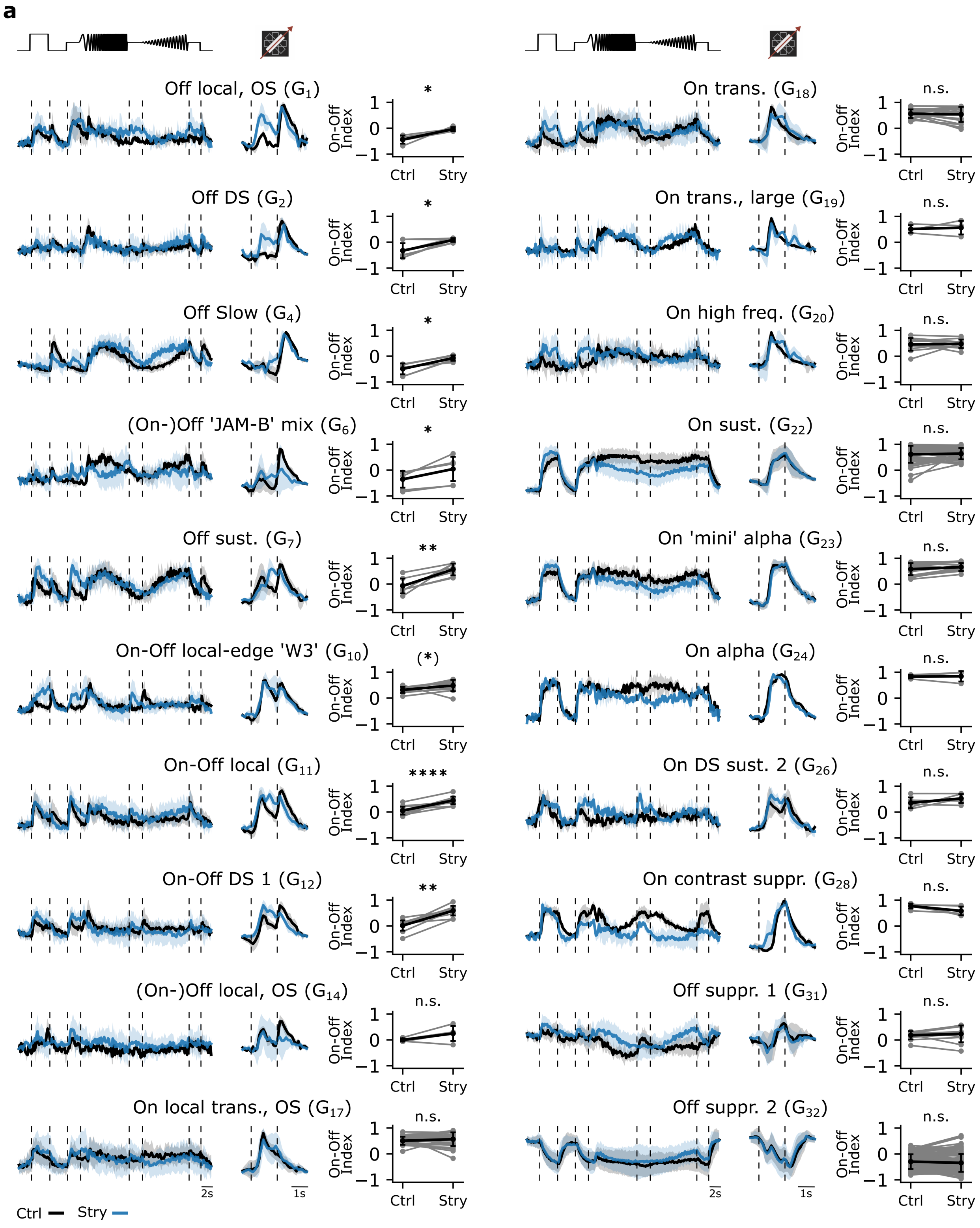


**Fig. S2. Testing the effects of strychnine on different RGC type responses. (a)** Mean $Ca^{2+}$ responses of sequentially recorded RGC types to the chirp (left) and moving bar (middle) to the Ctrl (black) and Strychnine (blue) conditions showing their corresponding On-Off indices for the Ctrl and Strychnine condition (right). RGC types with n>3 sequentially recorded cells per condition were included. ****: p<0.0001, ***: p<0.001, **: p<0.01, *: p<0.05, (*): p∼0.05; Paired T-Test.

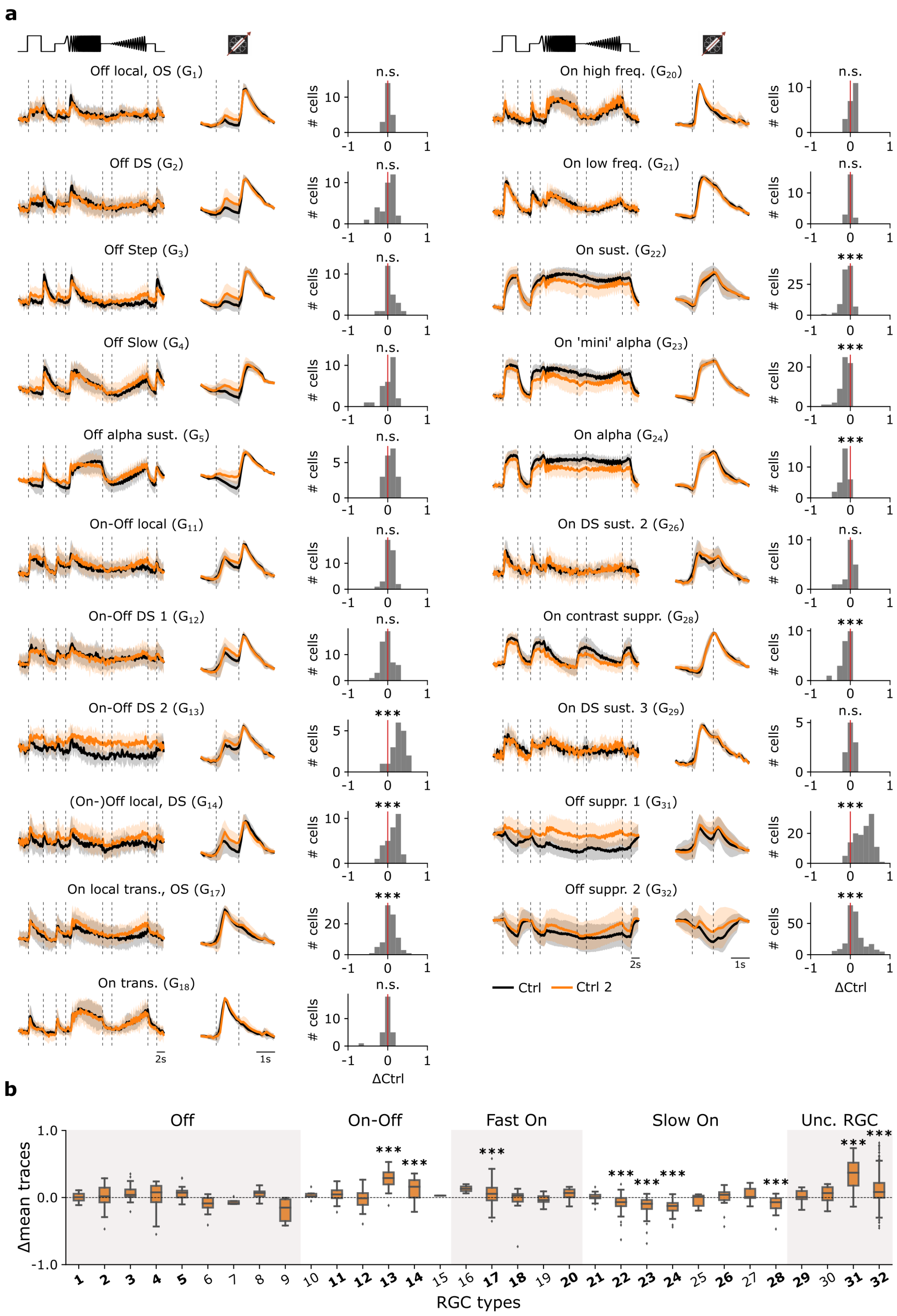


**Fig. S3. Adaptational, cell type-specific response changes without pharmacological pertubation. (a)** Mean $Ca^{2+}$ responses of sequentially recorded RGC types to the chirp (left) and moving bar (middle) to the Ctrl 1 (black) and Ctrl 2 (orange) conditions showing their trace differences (right). ***: p<0.001; One-sample T-Test. **(b)** Box plots of trace differences of all sequentially recorded cells of all RGC types from control-dataset (Ctrl 1 & Ctrl 2; orange). Bold numbers indicate RGC types with n>10 sequentially recorded cells per condition. Dashed line shows zero baselines, i.e., no difference between traces. Gray and white background blocks summarize the larger functional groups for better visualization (Off, On-Off, Fast On, Slow On, Uncertain RGCs). ***: p<0.001; One-sample T-Test.

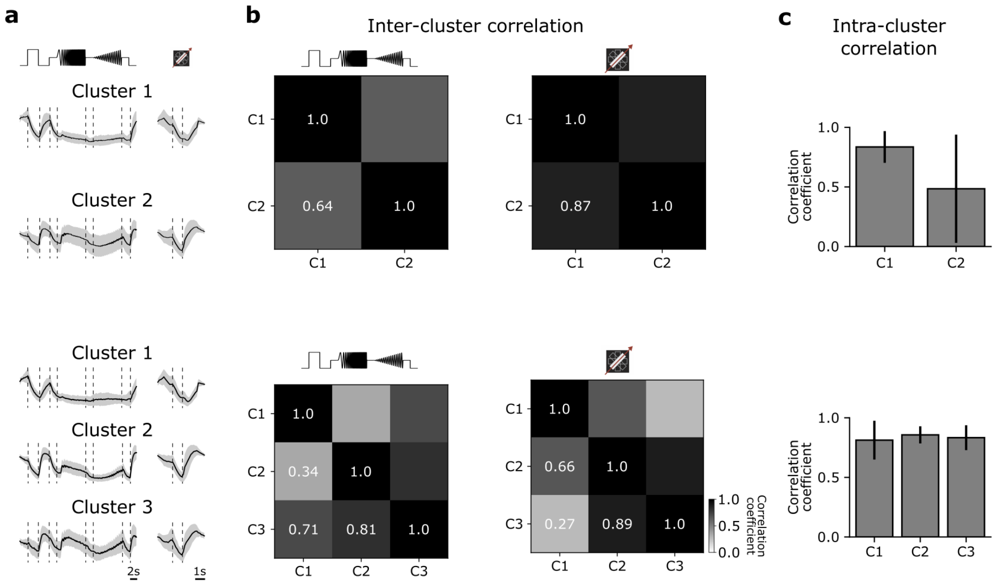


**Fig. S4. Further evaluation of the $G_{32}$ functional clustering. (a)** Top: Mean response to the chirp (left) and moving bar (right) using n=2 clusters for the functional clustering. Bottom: Mean response to the chirp (left) and moving bar (right) using n=3 clusters for the functional clustering. **(b)** Top: Inter-cluster correlation matrix between the mean responses of cluster 1 and cluster 2 of the chirp (left) and moving bar (right) responses. Bottom: Same as 'Top', but for n=3 clusters. **(c)** Top: Mean intra-cluster correlation for n=2 clusters with s.d. Bottom: Mean intra-cluster correlation for n=3 clusters with s.d.

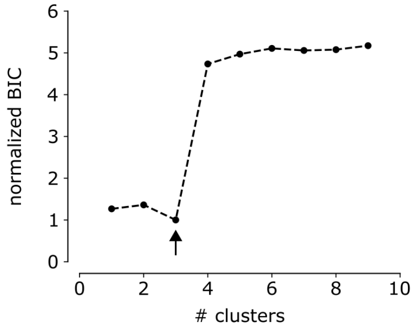


**Fig. S5. Bayesian Information Criterion as a function of number of clusters for $G_{32}$ in the MEA-dataset.** Bayesian Information Criterion (BIC) as function of number of clusters for the $G_{32}$ identified in the MEA-dataset. Arrow indicates the lowest BIC and the number of clusters to choose

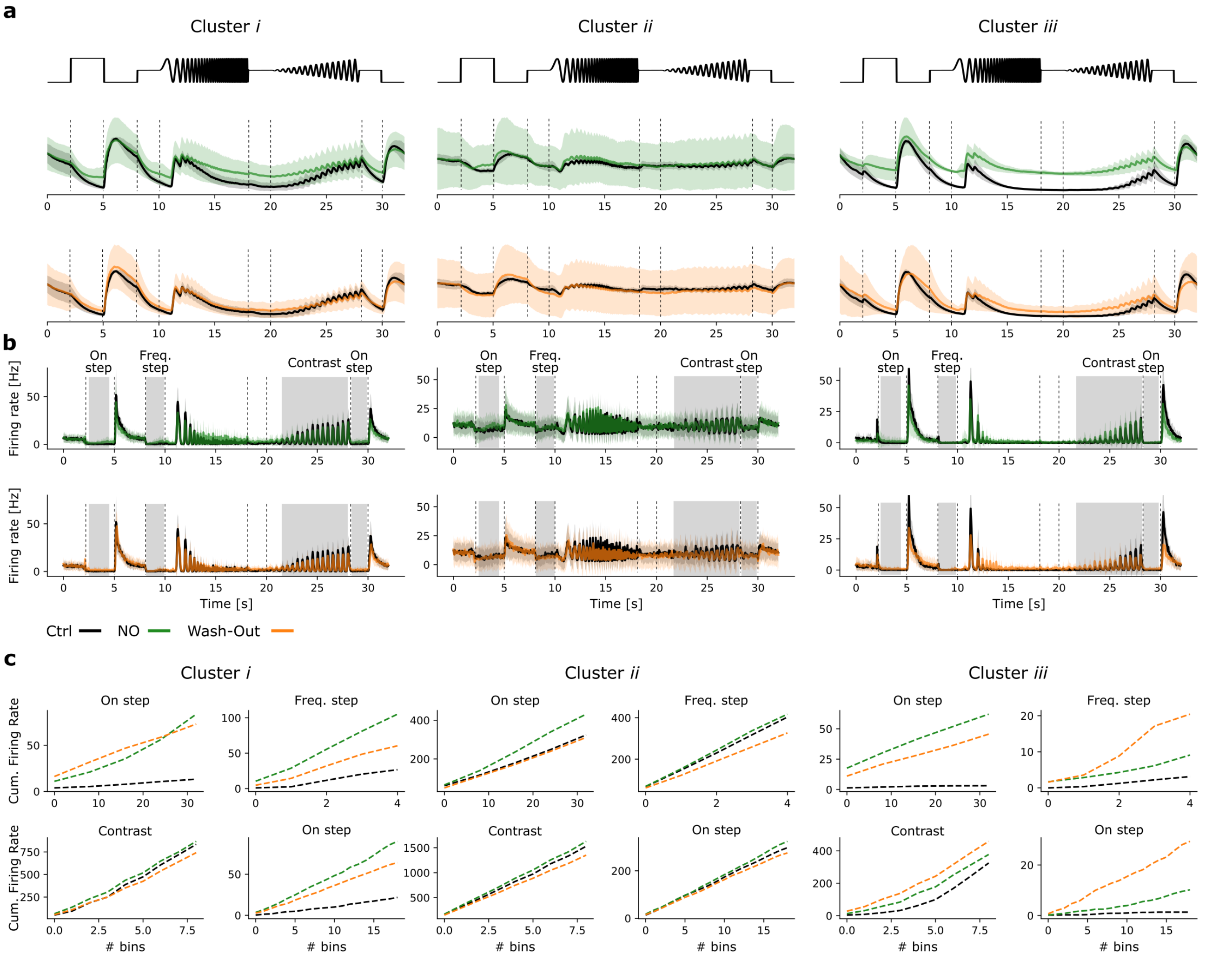


**Fig. S6. Pseudo-calcium and peristimulus time histograms of $G_{32}$ clusters. (a)** Pseudo-calcium traces of cluster *i*-*iii* (left to right) in response to the chirp under Ctrl (black) and NO (green; top) as well as Ctrl and Wash-out (orange; bottom) conditions. **(b)** Same clusters and conditions as (a), but peristimulus time histograms. **(c)** Cumulative firing rates of cluster *i*-*iii* (left to right) for the three conditions for four features (On step, frequency step, contrast, last on step).

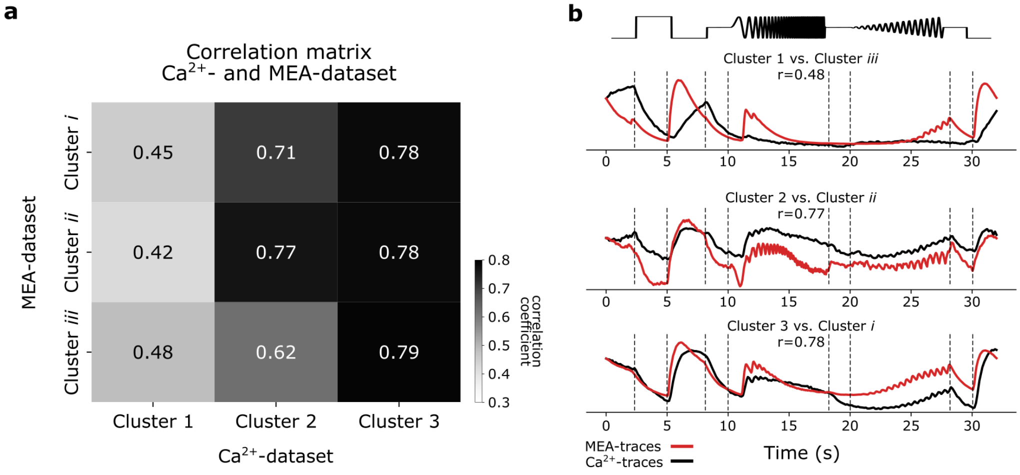


**Fig. S7. Correlating $G_{32}$ clusters of the $Ca^{2+}$-dataset and MEA-dataset. (a)** Correlation matrix showing the correlation coefficients between the pseudo-calcium $G_{32}$ cluster responses to the chirp and the $Ca^{2+}$-dataset $G_{32}$ clusters. Color bar indicates Pearson correlation coefficients. **(b)** Pseudo-calcium traces of $G_{32}$ clusters (red) overlaid with the correlated $Ca^{2+}$ $G_{32}$ cluster traces (black).
